# STRIDE: Signal Transfer and Aligned Ion-Peak Discrimination for Peptidoform Evidence for Consistent Quantification of Site-Localized Post-Translational Modifications in Large-Scale DIA-MS

**DOI:** 10.64898/2026.07.24.740615

**Authors:** Justin Cyril Sing, Shubham Gupta, Hannes Luc Röst

## Abstract

Data-independent acquisition (DIA) mass spectrometry has emerged as a powerful method for comprehensive post-translational modification profiling in proteomics, but faces challenges in consistently assigning peptidoforms to the same chromatographic peaks across multiple samples. This study introduces STRIDE to address this challenge by implementing dynamic programming alignment and propagating unique ion signature signals across runs. We benchmarked STRIDE using a synthetic phosphopeptide dilution series dataset, demonstrating an 11.3-fold reduction in false-negative rate compared to DIA-NN, from 68% with DIA-NN to 6% with STRIDE at 5% false-positive rate when requiring both correct site-localization and correct peak assignment. STRIDE also improved over the base OpenSWATH (IPF), reducing the false-negative rate from 14% to 6% for correct peak assignment. When applied to a phospho-enriched U2OS nocodazole dataset, our approach reduced missing values from 38.6% to 15.2% in a representative quantification matrix, increased highly reproducible phosphopeptide detections from 7,698 to 12,510 in nocodazole-treated samples, and identified 5,431 differentially abundant phosphopeptides compared to 2,588 with DIA-NN. This increased the number of enriched pathways recovered from differential phosphopeptide analysis, including pathways related to mitotic arrest, microtubule organization, chromatin regulation, RNA processing and kinase signalling. Lastly, we analyzed acetyllysine and glycopeptide-enriched DIA datasets, where STRIDE improved complete replicate detection, reduced missing quantification values, and increased site-specific glycan reproducibility. Together, these results demonstrate that STRIDE improves peptidoform proteomics through reduced false-negative rates, consistent peak assignment, and more complete quantification across runs.

## Introduction

Post-translational modifications (PTMs) play a crucial role in biology, contributing to the functional diversity of the proteome and allowing for a greater range of functions than those encoded by the genome.^1–3^ PTMs help regulate protein activity, localization, stability, and interactions by covalently modifying amino acid side chains or protein termini, altering the physicochemical and biochemical properties of proteins.^1–3^ PTMs, such as phosphorylation, glycosylation, acetylation, sumoylation, and many others, can act as molecular switches, allowing cells to respond to rapid environmental changes and internal signals.^3,4^ They are involved in numerous biological processes, including cell signalling, protein degradation, and cellular metabolism.^3,5^ Hence, understanding PTMs is an important aspect of systems biology for deciphering complex cellular networks, elucidating disease mechanisms, and developing targeted therapies for fields like cancer research.^3,5^ More than 680 types of PTMs have been identified to date, and current PTM resources contain hundreds of thousands of experimentally observed modification sites; for example, PhosphoSitePlus reports over 494,000 non-redundant PTM sites.^6–9^ The large diversity of modifiable amino acids and combinatorial possibilities makes identification and localization of PTMs challenging.^2^

Liquid chromatography-tandem mass spectrometry (LC-MS/MS) has become the major high-throughput approach for detecting and quantifying PTMs. In bottom-up LC-MS/MS proteomics, proteins are enzymatically digested into peptides and analyzed using acquisition strategies such as discovery-based data-dependent acquisition (DDA)^10,11^, targeted selected or parallel reaction monitoring (SRM^12^ or PRM^13^), and data-independent acquisition (DIA)^11,14–18^. SWATH-MS, a variant of DIA, systematically fragments all precursors within predefined mass-to-charge (m/z) isolation windows (typically 25 m/z) that iteratively change across a mass range (i.e., 400-1200 m/z)^19^. SWATH-MS combines the identification breadth of DDA and the reproducibility of PRM, making it suitable for investigating the PTM landscape.^10,14^

DIA methods are better suited than DDA for consistent coverage and quantification of peptidoforms across large cohorts. ^17^ However, applying DIA to modified peptides introduces a different set of challenges, specifically with isomeric PTMs: peptidoforms that share the same backbone sequence and the same precursor mass but differ only in the site of modification. The only information that separates them is a small subset of site-specific fragment ions. Rosenberger et al. demonstrated this with a hierarchical-Bayesian framework that propagates the confidence of site-localization detection at the precursor-level, transition-level, and peptidoform-level, showing that DIA can reliably identify and site-localize PTMs with high confidence.^14^

However, due to the high similarity of chromatographic peak groups originating from different peptidoforms of the same peptide sequence, accurate quantification across large-scale experiments remains highly challenging due to incorrect or missing peak-group assignment for isomeric peptidoforms. In some samples, the inference algorithm can assign one peptidoform to a first eluting peak; while in others the same peptidoform is assigned to a secondary eluting peak or missed entirely. These inconsistent assignments are detrimental to downstream biological analyses. Importantly, this is not an edge case; it is the expected outcome in any large-scale DIA experiment where isomeric PTMs co-elute closely, and to our knowledge no current tool systematically addresses it. To address this challenge, we developed STRIDE, an cross-run peptidoform-inference workflow that maps extracted ion chromatographic peak-groups and propagates unique ion signature (UIS) signals across runs to boost inference of peptidoforms.

Gupta. *et al.* previously demonstrated a novel dynamic programming alignment method for aligning extracted ion chromatograms for DIA-MS proteomics data.^20,21^ We extend this method for generating precursor peak-group mappings across runs for all potential peak-groups, and implement aligned peak-group scoring for computing error estimates for the quality of mapped aligned peak-groups, as well as scoring for the aligned identification transitions. Drawing from the principles of inference of peptidoforms^14^ and the quantification recovery of DIAlignR^20,21^, we developed a signal transfer and aligned ion-peak discrimination for peptidoform evidence (STRIDE) method to bridge the gap between confident site-localization and consistent quantification. We benchmarked STRIDE on a “ground truth” synthetic phosphopeptide dataset with manual annotations, a biologically complex phosphopeptide-enriched dataset as well as application to other types of PTMs (acetyllysine, glycosylation).

## Results

### Cross run inference for consistent identification and quantification of peptidoforms in large scale data-independent mass spectrometry experiments

The STRIDE method was developed to consistently site-localize PTMs across experimental data-independent acquisition (DIA) mass spectrometry runs.

In our workflow, OpenSWATH first performs peak picking for each precursor to identify candidate peak-groups in each run (Fig. 1A). STRIDE then aligns the extracted ion chromatograms for the same precursor across runs to establish correspondence between peak-groups. By default, this alignment is driven by the detecting (empirical) fragment-ion traces, which are collapsed into a precursor-specific total ion chromatogram (TIC) for each run. These TICs are smoothed, normalized, and aligned across runs using an initial fast Fourier transform (FFT)-based retention-time shift followed by dynamic time warping refinement. The resulting alignment path defines a retention-time mapping between the reference run and each query run, which is then used to project reference peak-group apices and boundaries into the other runs (Supp. Fig. 1).

**Figure 1.**
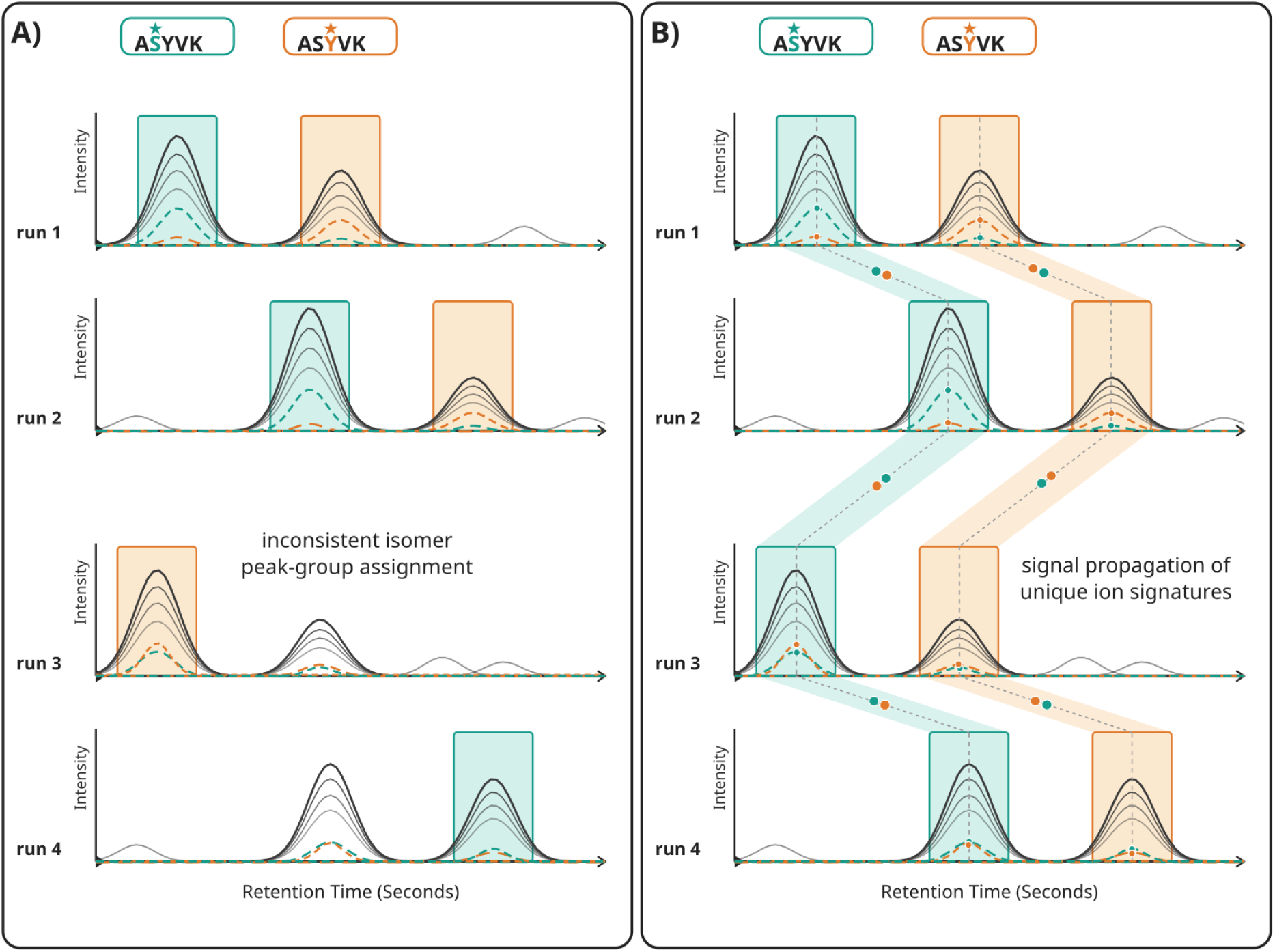
Cross-Run Inference of Peptidoforms. This schematic illustrates the challenge of consistently quantifying peptidoforms that share the same peptide backbone sequence, ASYVK, but differ in phosphorylation-site localization across runs. **A)** The left panel shows an example case where in two runs the first peptidoform gets assigned to the first eluting peak, and the second peptidoform gets assigned to the second eluting peak. However, in the last two runs, either a peritoform is missed from identification or is assigned to an incorrect peak by the current state-of-the-art inference algorithm. **B)** The second panel demonstrates the cross-run inference algorithm which aligns the peak-groups across runs and then propagates the unique ion signature signals across runs to boost the inference of peptidoforms for confident and consistent peptidoform quantification across runs.

For each projected reference peak-group, STRIDE evaluates candidate peak-groups already detected in the corresponding query run. Rather than selecting the nearest peak by retention time alone, candidate mappings are scored using a combination of retention-time agreement, peak-boundary overlap, peak-width similarity, intensity similarity, and existing OpenSWATH peak-group scoring metadata. STRIDE then performs a one-to-one assignment so that each query peak-group can only be mapped to a single reference peak-group (Supp. Fig. 1). Each selected mapping is assigned an additional confidence score that reflects the strength of the selected candidate, its separation from nearby alternatives, the local consistency of the retention-time mapping, and the ambiguity of the local candidate space (Supp. Fig. 3). A more detailed description of the alignment, candidate assignment, and mapping confidence calculation is provided in the algorithm overview in the supplement.

The aligned peak mappings are then used during inference of peptidoforms. Identifying transitions are not used to force a peptidoform assignment from one run to another. Instead, STRIDE uses confident cross-run peak mappings to transfer already-supported transition evidence between aligned peak-groups. The peptidoform posterior probabilities are then recomputed using the augmented evidence set, so transferred evidence contributes probabilistically rather than acting as deterministic label copying (Fig. 1B, Supp. Fig. 2). This allows unique and shared ion evidence observed in one run to support the same aligned peak-group in other runs, while restricting propagation to mappings and transition evidence that pass confidence-based filters.

### Benchmarking on Synthetic Phosphopeptide Dilution Series

To assess the performance of STRIDE, we analyzed a synthetic phosphopeptide dataset previously published by Rosenberger *et al.*^14^. Briefly, the experiment used 579 synthetic, heavy-isotope-labeled phosphopeptides from *S. cerevisiae* proteins spiked into a human cell line (HEK-293) background as a 13-step dilution series to create a reference dataset for benchmarking the performance of the IPF method, specifically focusing on the ability to correctly site-localize phosphorylation and to test the detection limit of low-abundance phosphopeptides.^14^ For analysis, we used DIA-NN as the primary comparison method because it is a leading DIA analysis platform and is widely used in the field, including PTM-focused DIA workflows. DIA-NN also includes dedicated peptidoform scoring and match-between-run reanalysis support, making it a strong benchmark for evaluating the performance of STRIDE. For this dataset, DIA-NN was run using the peptidoform analysis settings recommended in the DIA-NN documentation (see methods for specific parameters).

For this dilution series dataset, there is a greater likelihood that peptidoform peak assignments will vary across runs or that identifications will be missed, especially at higher dilution steps and lower peptide concentrations, due to differences in signal quality. For example, the extracted ion chromatograms for phosphorylated peptide EST(Phospho)AEPDSLSR demonstrate this for DIA-NN, where DIA-NN consistently selects the wrong secondary peak in four out of six runs, while not assigning any peaks to the peptidoform in the other two runs (Fig. 2A, Supp. Fig. 4). When applying STRIDE, the peptidoform peak boundaries are assigned more consistently across dilution steps and better match the manually annotated Skyline peak boundary regions (Fig. 2A). To further investigate the performance not just in correct site-localization of the peptidoform, but also for correct peak assignment, we manually annotated 173 peptidoform precursors across the 13 dilution steps in Skyline^22^. When considering both the correct site-localized peptidoform and the correct peak assignment, STRIDE achieves a 94% true positive rate (TPR) at 5% false-positive rate (FPR), compared to 32% TPR for DIA-NN (Fig. 2B). Across dilution steps, STRIDE consistently detects more true spiked-in phosphopeptides than DIA-NN at 5% peptidoform false discovery rate (FDR), with a larger gain at lower peptide concentrations while maintaining a lower number of false peptidoforms across dilution series (Fig. 2C). Furthermore, examining the quantification performance, DIA-NN’s quantification on average is higher than the expected ground truth, with greater variation in intensity across the dilution series (Supp. Fig. 6C).

**Figure 2.**
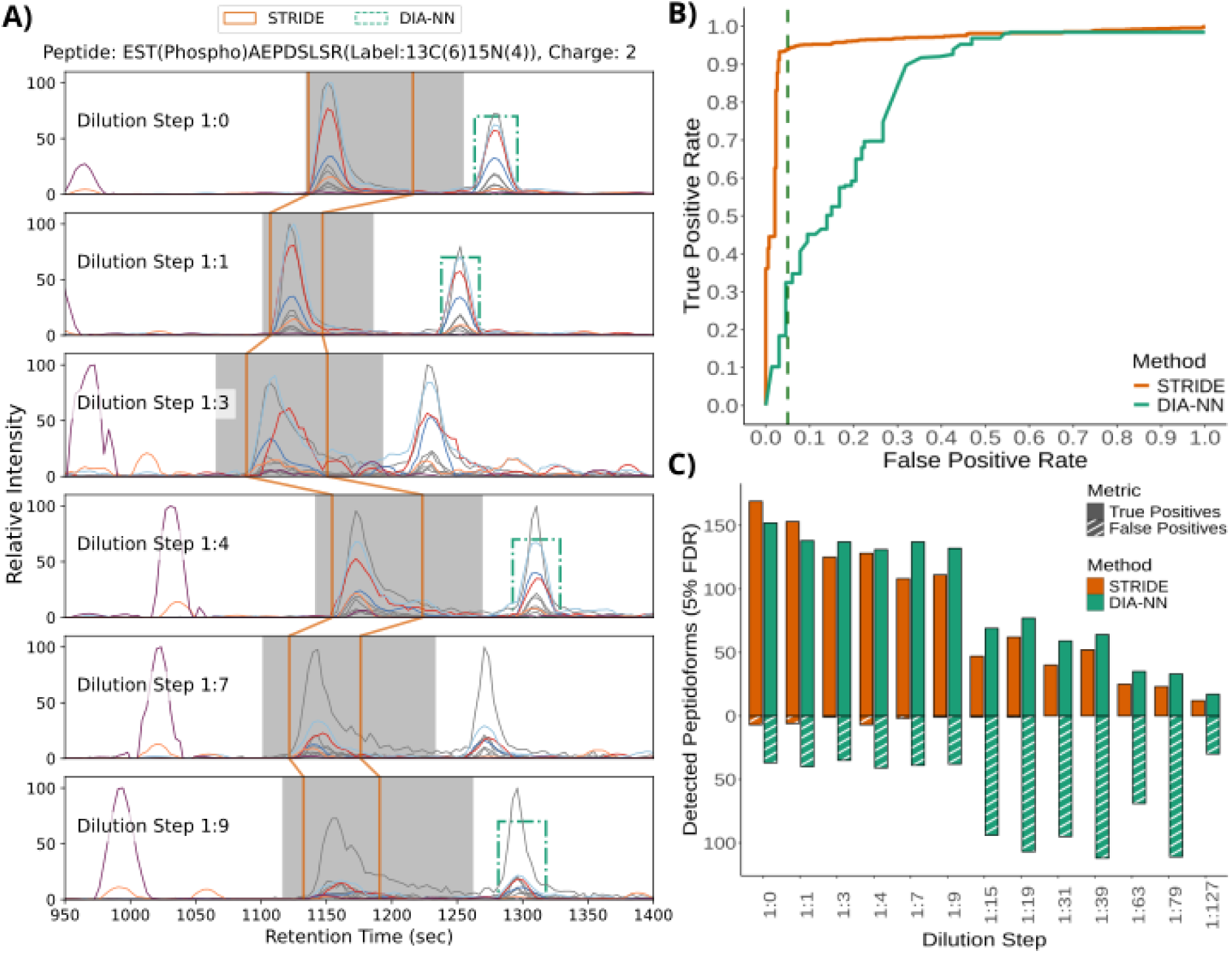
Improved Accuracy of Consistent Quantification of Peptidoforms on a Synthetic Phosphopeptide Dataset. **A)** Example extracted ion chromatograms for phosphorylated peptide EST(Phospho)AEPDSLSR across the first 6 dilution steps. The grey traces represent the empirical peak-group detecting transitions, and the colored traces represent unique-ion signatures for the b3 and y9 ions. The grey shaded region represents the manually annotated region from Skyline, the green dashed rectangles represent the peak boundaries assigned by DIA-NN at 5% peptidoform false discovery rate (FDR), and the orange boundaries connected across the runs represent peak boundaries assigned by STRIDE at 5% FDR. **B)** The receiver operating characteristic (ROC) curve shows the identification performance comparing DIA-NN and STRIDE. The solid lines show the performance of correctly identifying the right peptidoform as well as correctly assigning it to the correct peak (from manual annotations). DIA-NN at 5% FPR achieves 32% TPR and STRIDE achieves 94% TPR at 5% FPR. **C)** The bar plot shows the number of identifications per dilution step at a peptidoform level FDR at 5%. The orange bars represent STRIDE and the green bars represent DIA-NN. The solid bars represent the true positives, and the striped bars represent the false positives.

For comparison, we also evaluated the standard IPF workflow. With IPF, looking at the same example extracted ion chromatogram for EST(Phospho)AEPDSLSR demonstrates the issue of inconsistent peptidoform-peak assignment. The third and sixth dilution steps select the wrong secondary peak, whereas runs one, two, four and five select the first peak (Supp. Fig. 4). Looking at the ROC curve, STRIDE improves the identification performance compared to IPF, achieving 94% TPR compared to 87% TPR for IPF at a 5% FPR when assessing only correct peptidoform identification (Supp. Fig. 6A). When considering both correct site-localized peptidoform identification and correct peak assignment, STRIDE maintains 94% TPR, while IPF reaches 86% TPR at 5% FPR (Supp. Fig 5A). This can be further seen for the RGS(Phospho)VYHPLNIVQADAVR peptidoform, where STRIDE is able to recover peaks in the lower concentration dilution steps that IPF and DIA-NN were not confidently able to assign to the peptidoform (Supp. Fig. 5).

These results demonstrate that STRIDE improves both sensitivity and consistency in peptidoform identification and quantification, particularly at challenging lower peptide concentrations. On this benchmark synthetic phosphopeptide dataset, STRIDE outperforms DIA-NN in correctly assigning site-localized peptidoforms compared to the manually annotated peak, while also improving peak assignment consistency and recovery of lower-abundance peptidoforms compared to IPF.

### Improving the Quantification Matrix Completeness in a U2OS Phospho-enriched Dataset

We next investigated whether the benefits of STRIDE extend beyond benchmark data to real biological datasets. We thus analysed a phospho-enriched U2OS dataset previously published in Rosenberger *et al.*^14^ which contains phosphopeptide-enriched samples from human U2OS cells treated with nocodazole versus untreated controls across ten biological replicates. Nocodazole disrupts spindle microtubules and induces mitotic arrest, providing a model system with expected changes in phosphorylation-associated signaling.

To evaluate whether STRIDE improves the phosphopeptide quantification in a more complex biological dataset, we applied DIA-NN and STRIDE to the phospho-enriched U2OS dataset. Visual inspection of matched phosphopeptide-by-run quantification matrices showed that DIA-NN contained more missing values across both control and treated samples, whereas STRIDE produced a more complete matrix with stronger across-run consistency (Fig. 3A). In the representative set of 100 random sampled phosphopeptides, missing values decreased from 38.6% with DIA-NN to 15.2% with STRIDE, while the mean number of quantified runs per phosphopeptide increased from 12.3 to 16.9 (Fig. 3A). This improvement was also observed in the full quantification matrix for intersecting phosphopeptides, where the STRIDE matrix contained fewer missing values while preserving comparable phosphopeptide intensity distributions across samples (Supp. Fig. 8A,C).

**Figure 3.**
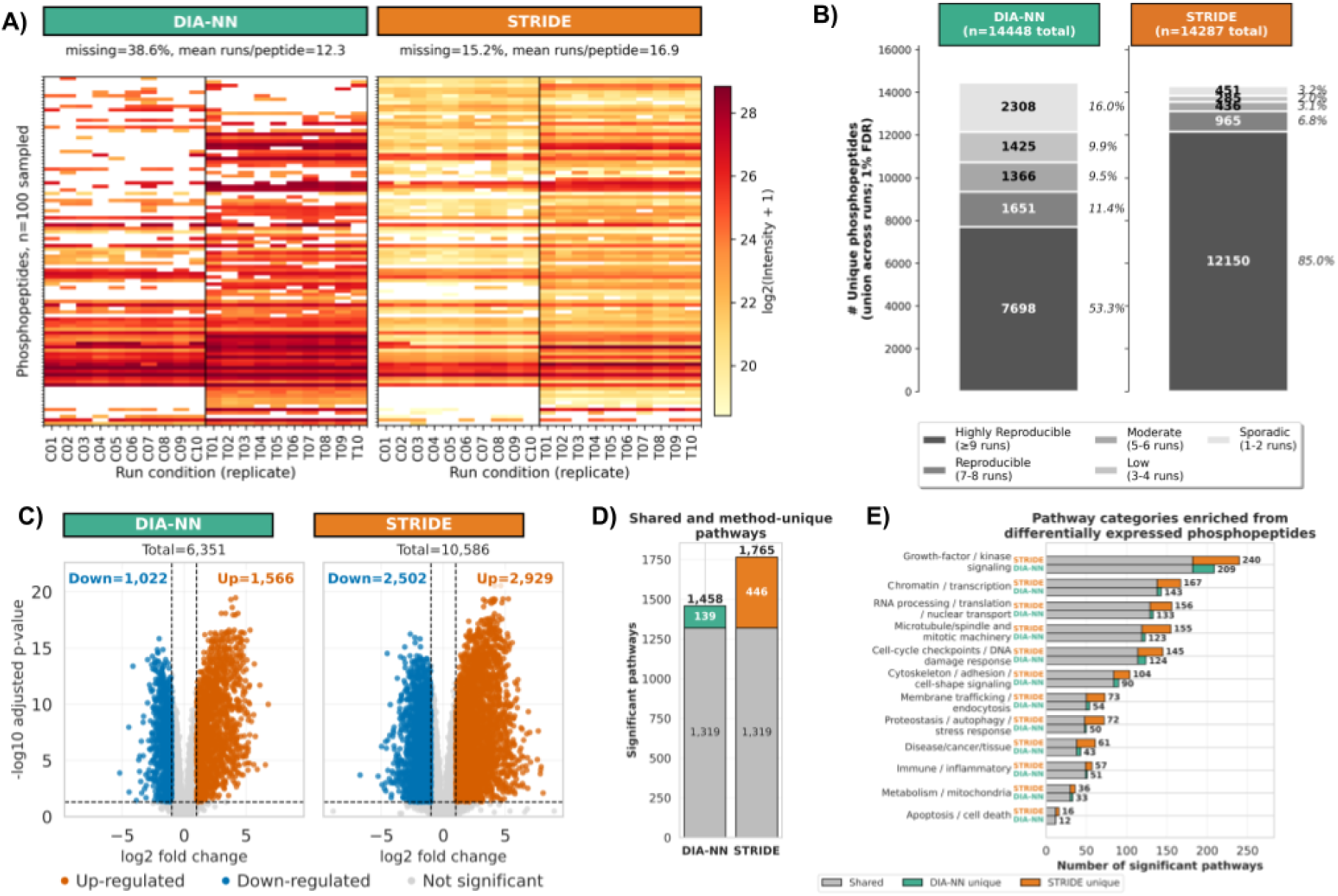
STRIDE improves phosphopeptide identification consistency and downstream pathway analysis in a U2OS phospho-enriched nocodazole dataset. **A)** Quantification matrices of 100 randomly sampled phosphopeptides across 10 control and 10 nocodazole-treated samples for DIA-NN and STRIDE. Rows represent phosphopeptides and columns represent runs ordered by condition and replicate. Color indicates log2(Intensity + 1), and white entries indicate missing values. The percentage of missing values and mean number of quantified runs per peptide are shown above each matrix. **B)** Reproducibility of unique phosphopeptide identifications across runs at 1% FDR. Stacked bars show the number of phosphopeptides detected across the indicated number of runs, ranging from highly reproducible (>=9 runs) to sporadic (1-2 runs). **C)** Volcano plots of differentially abundant phosphopeptides between nocodazole-treated and control samples for DIA-NN and STRIDE. Up-regulated phosphopeptides are shown in orange, down-regulated phosphopeptides in blue, and non-significant phosphopeptides in grey using an FDR threshold of 0.05 and an absolute log2 fold-change threshold greater than 1. **D)** Shared and method-unique significant pathway enrichments identified from differentially abundant phosphopeptides. Grey indicates pathways shared by both methods, green indicates DIA-NN-unique pathways, and orange indicates STRIDE-unique pathways. **E)** Category-level summary of significant pathway enrichments from differentially abundant phosphopeptides. Bars show shared and method-unique pathways within each biological category.

We next assessed whether improved quantification translated into more reproducible phosphopeptide detection across runs. Although the total number of unique phosphopeptides detected in nocodazole-treated samples was similar between DIA-NN and STRIDE, the distribution of identifications across runs differed substantially (Fig. 3B). STRIDE detected 12,510 phosphopeptides in at least 9 of 10 treated samples, compared with 7,698 for DIA-NN, and reduced the number of sporadically detected phosphopeptides observed in only 1–2 runs from 2,308 to 451 (Fig. 3B). Consistent with this, per-replicate identification counts were higher for STRIDE across both control and nocodazole-treated samples, and run-sampling analyses showed that STRIDE retained a larger fraction of phosphopeptides as the number of runs increased (Supp. Fig. 7A,C). The narrow sampling intervals observed for STRIDE further indicated reduced run-to-run variability in identification consistency (Supp. Fig. 7D). Across all samples and within each condition separately, STRIDE also increased the number and fraction of phosphopeptides quantified in every run (Supp. Fig. 7G).

We then evaluated how these changes affected downstream differential expression analysis. Using an adjusted p-value threshold of 0.05 and absolute log2 fold-change greater than 1, DIA-NN identified 1,566 up-regulated and 1,022 down-regulated phosphopeptides from 6,351 tested phosphopeptides, whereas STRIDE identified 2,929 up-regulated and 2,502 down-regulated phosphopeptides from 10,586 tested phosphopeptides (Fig. 3C). Additional comparisons of differentially abundant phosphopeptides and their mapped proteins showed both shared and method-specific differential signals, including proteins with increased and decreased phosphorylation detected by each method (Supp. Fig. 9A,B). When differential expression analysis was repeated while requiring quantification in increasing numbers of replicates, STRIDE retained more differentially abundant phosphopeptides across strict replicate requirements (Supp. Fig. 9C), consistent with the improved run completeness observed in the quantification-level analyses.

Finally, we assessed whether the increased number of differentially abundant phosphopeptides translated into pathway-level information. Pathway enrichment analysis identified 1,319 pathways shared between DIA-NN and STRIDE, 139 pathways unique to DIA-NN, and 446 pathways unique to STRIDE (Fig. 3D). Categorizing enriched pathways by biological theme showed that STRIDE increased pathway coverage across several categories relevant to nocodazole perturbation, including microtubule/spindle and mitotic machinery, cell-cycle checkpoints and DNA damage response, chromatin/transcription, RNA processing and nuclear transport, and growth-factor or kinase signalling (Fig. 3E). Mirrored pathway-enrichment plots further showed the shared strength among pathways detected by both methods (Supp. Fig. 9D,E).

To place the pathway-level results within the expected nocodazole response, we summarized the differential phosphoprotein and phosphosite signals as a working model of checkpoint-maintained mitotic arrest (Fig. 4).^23^ Nocodazole-induced spindle perturbation activates spindle-checkpoint regulation of APC/C-CDC20, and inhibition of APC/C-CD20 maintains mitotic arrest. Differential phosphorylation associated with BUB1, BUB3 and MAD2L1BP connected the recovered signal to spindle-checkpoint regulation.^24–26^ These associations do not establish the direction or functional consequence of measured phosphorylation changes.

**Figure 4.**
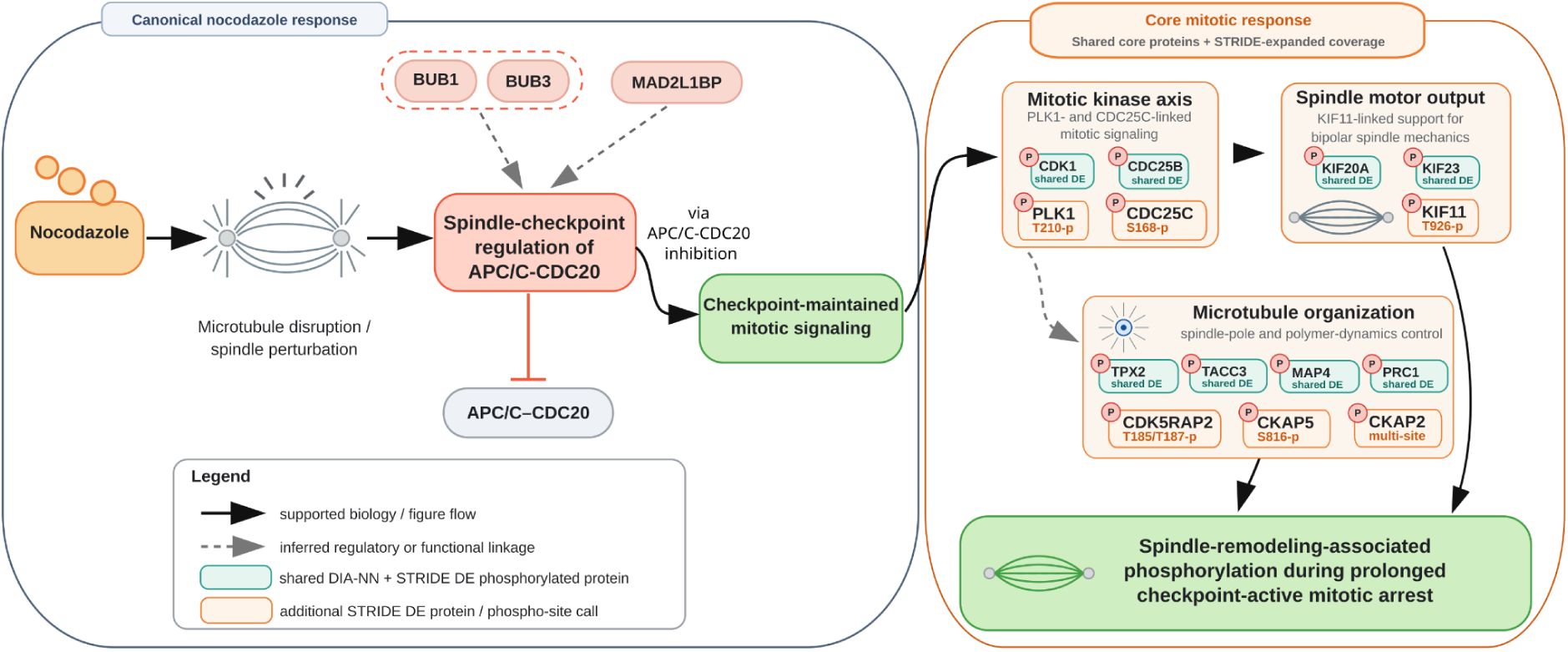
Shared and STRIDE-expanded phosphorylation signals support a checkpoint-maintained mitotic state in nocodazole-treated U2OS cells. The left panel summarizes the canonical nocodazole response. Nocodazole disrupts spindle microtubules and produces spindle perturbation, leading to spindle-checkpoint regulation of APC/C-CDC20. Inhibition of APC/C-CDC20 maintains mitotic arrest. BUB1 and BUB3 are grouped as spindle-checkpoint regulators, while MAD2L1BP is shown separately as an additional checkpoint-associated protein. Dashed connections indicate inferred regulatory or functional associations and do not assign a functional consequence to the measured phosphorylation changes. The right panel summarizes the differential phosphoprotein and phosphosite signals within mitotic kinase, spindle motor, and microtubule-organization modules. Teal outlines indicate shared DIA-NN and STRIDE proteins mapped from differentially abundant phosphopeptides, and orange outlines indicate additional STRIDE differential protein or phosphosite calls. Solid black arrows indicate supported biological or figure flow, the red inhibitory line indicates APC/C-CDC20 inhibition, and dashed grey arrows indicate inferred regulatory or functional linkages.

The broader differential signal contained a shared core mitotic response together with additional STRIDE-expanded proteins mapped from differentially abundant phosphopeptides and site-level coverage (Fig. 4). Shared DIA-NN and STRIDE differential signals include proteins linked to the mitotic kinase axis, spindle motor output, and microtubule organization, including CDK1, CDC25B, KIF20A, KIF23, TPX2, TACC3, MAP4 and PRC1.^27–34^ STRIDE added differential phosphosite calls involving PLK1 (T210), CDC25C (S168), KIF11 (T926), CDK5RAP2 (T185 and T187), CKAP5 (S816), and CKAP2 (S534 and S597).^35–39^ PLK1 (T210), CDC25C (S168), KIF11 (T926) are previously characterised regulatory phosphorylation sites^40–42^, while the detected CDK5RAP2, CKAP5, and CKAP2 sites provide additional site-level coverage of protein with established roles in spindle and microtubule organization^36–39^. These additional calls mapped to the same mitotic processes recovered by both methods, supporting a more complete phosphoprotein- and site-level view of spindle-remodeling-associated phosphorylation during prolonged checkpoint-active mitotic arrest.

Within the expanded phosphopeptide results, CKAP2 provided a multi-site example of the additional site-level information recovered by STRIDE (Supp. Fig. 10). Detected CKAP2 phosphopeptides mapped to S304, S534, T579/T582, T597, S614 and T683 across the protein sequence. Several phosphopeptides increased in nocodazole-treated samples, including S534- and T597-containing phosphopeptides called differentially abundant by STRIDE but not DIA-NN. STRIDE also quantified CKAP2 phosphopeptides more completely across individual runs, making the coordinated multi-site pattern more apparent.

Together, these results show that STRIDE improves phosphopeptide quantification completeness and reproducibility, which increases the number of phosphopeptides available for differential expression analysis and expands the downstream pathway-level information recovered from the U2OS nocodazole experiment.

### Evaluating Performance on Other Types of Post-Translational Modifications

To assess whether the STRIDE provides similar benefits to other types of PTMs, we analyzed a previously published acetyllysine-enriched peptide dataset by Meyer, J. *et al.*^16^ and a glycopeptide-enriched dataset by Yang, Y. *et al.*^43^ Briefly, for the first dataset, acetylated-enriched peptides were acquired from mouse liver tissue samples, with triplicate samples of 0.5-fold volume (termed halfDIA hereafter) injected, and another set of triplicate runs acquired with 1.0-fold volume (termed fullDIA hereafter) of enriched acetylated peptides^16^. The purpose of this experimental setup is to assess the quantification performance between the halfDIA and fullDIA samples. The second dataset, consisting of glycosylated-enriched peptides from a pooled mixture of 40 human serum samples, was acquired and processed in three replicate runs.^43^

In the acetyllysine dataset, STRIDE increased the number of unique acetyllysine peptides detected in at least one replicate from 921 (detected by DIA-NN) to 1,319 in the halfDIA condition, a 1.43-fold gain. In the fullDIA condition, STRIDE increased the total from 1,080 to 1,390 peptides, a 1.29-fold gain. The improvement was larger when considering complete observations across replicate runs. In the halfDIA condition, the number of acetyllysine peptides detected in all three replicates increased from 668 to 1,106, corresponding to a 1.66-fold increase and an increase in quantification matrix completeness from 72.% to 83.9%. In the fullDIA condition, complete identifications increased from 652 to 1,115, corresponding to a 1.71-fold increase and an increase in completeness from 60.4% to 80.2% (Fig. 5A). Across all six runs, STRIDE doubled the number of acetyllysine peptides detected in every run, from 507 to 1,015, and increased the complete fraction from 44.6% to 72.6% (Supp. Fig. 12C). This also reduced missing quantification values by 44.6% in the halfDIA condition and 51.9% in the fullDIA condition (Fig. 5B).

**Figure 5.**
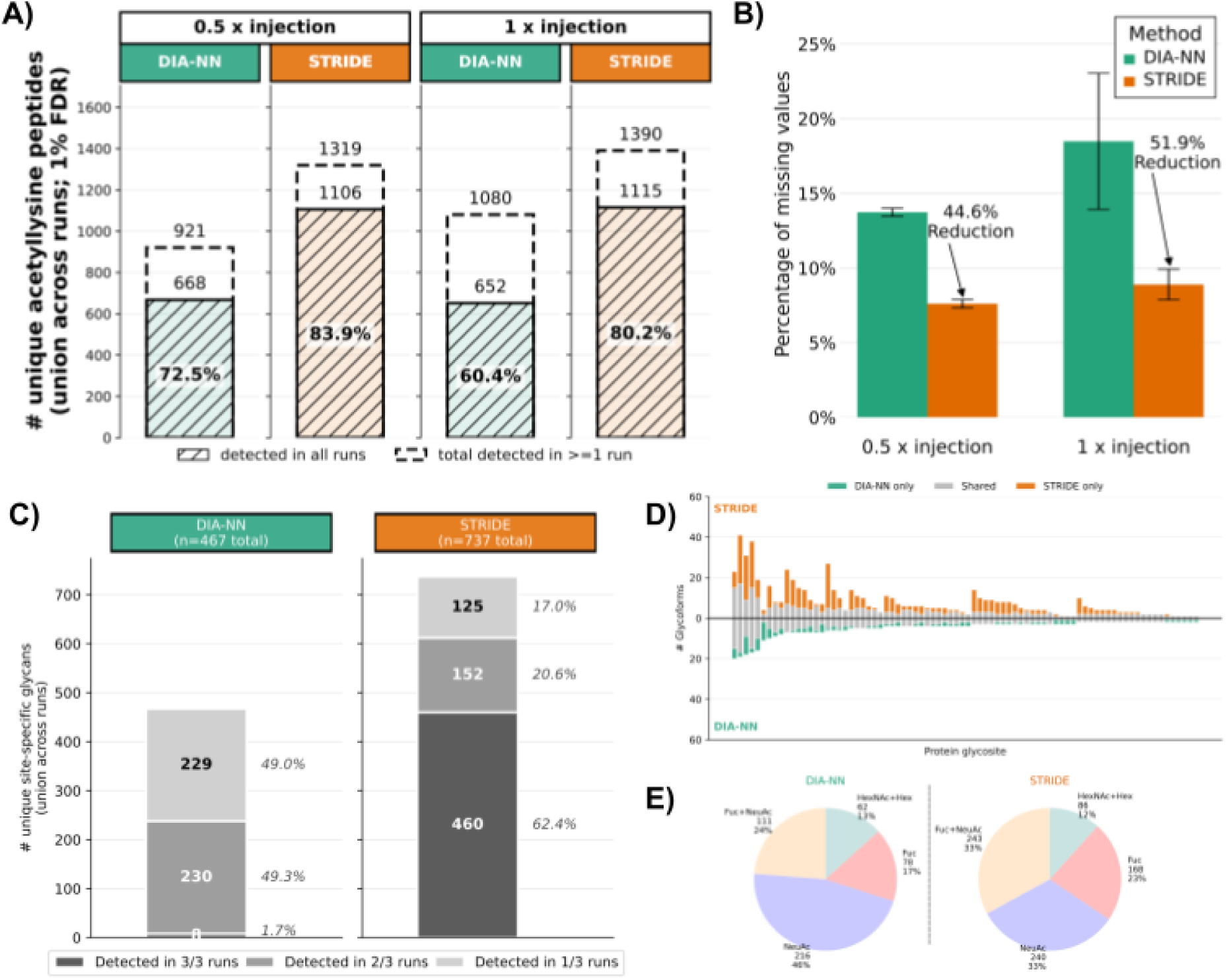
STRIDE improves reproducibility and data completeness for acetyllysine and glycopeptide DIA analyses. **A)** Complete versus total acetyllysine peptide identifications in the acetyllysine-enriched DIA dataset. Identifications were filtered at 1% site-localization FDR and 1% global peptide FDR. Dashed bars show the total number of unique acetyllysine peptides detected in at least one run, while hatched bars show the subset detected in all three replicate runs for each injection condition. **B)** Mean percentage of missing quantification values across replicate runs for DIA-NN and STRIDE. Error bars show the variation across runs, and arrows indicate the reduction in missing values obtained with STRIDE. **C)** Reproducibility of site-specific glycan identifications in the human serum N-glycopeptide dataset. Stacked bars show the number and percentage of site-specific glycans detected in 3/3, 2/3 or 1/3 replicate runs. **D)** Mirror plot comparing the number of glycoforms assigned to each protein glycosite by DIA-NN and STRIDE. Shared glycoforms are shown in grey, DIA-NN-only glycoforms in green, and STRIDE-only glycoforms in orange. **E)** Distribution of glycoform composition classes identified by DIA-NN and STRIDE in the human serum N-glycopeptide dataset.

The additional acetyllysine identifications were not limited to a small method-specific tail. At 1% peptidoform FDR, DIA-NN and STRIDE shared 1,036 peptide-level identifications, but STRIDE added 303 unique peptide-level identifications compared with 69 DIA-NN-only identifications. At the acetyllysine level, the same pattern was observed, with 1,063 shared identifications, 336 STRIDE-only identifications, and 73 DIA-NN-only identifications (Supp. Fig. 11C). The quantification heatmap showed that STRIDE filled in missing values across replicate runs rather than only adding isolated single-run observations, and the log2 fold-change distributions for fullDIA relative to halfDIA remained centred near the expected value of 1 (Supp. Fig. 12A,F-G).

In the human serum N-glycopeptide dataset, STRIDE produced a larger effect on reproducibility. At the site-specific glycan level, the number of unique identifications increased from 467 with DIA-NN to 737 with STRIDE, an increase of 270 site-specific glycans or 1.58-fold. More importantly, the number detected in all three replicates increased from 8 site-specific glycans with DIA-NN to 460 with STRIDE, increasing the complete fraction from 1.7% to 62.4% (Supp. Fig. 13E,H). This suggested that STRIDE mainly improves the consistency of glycoforms across runs, rather than simply increasing the total number of reported sites.

The missing-value analysis supports this interpretation. STRIDE reduced missing quantification values by 63.1% at the glycopeptide level, 63% at the site-specific glycan level, and 81.5$ at the protein glycosite level (Supp. Fig. 13J-L). In the quantification heatmaps, DIA-NN had 46.8% missing values for site-specific glycans and 36.6% missing values for protein glycosites, whereas STRIDE reduced these to 11.0% and 4.8%, respectively (Supp. Fig. 14A-B). Quantification variability also improved at the more biologically summarized levels. The median glycopeptide-level coefficient of variation (CV) was similar between methods, 14.2% for DIA-NN and 14.5% for STRIDE, but the median site-specific glycan CV decreased from 17.4% to 13.5%, and the median protein glycosite CV decreased from 31.7% to 15.9% (Supp. Fig. 14F-H).

The expanded site-specific glycan identifications covered all the major glycoform composition classes. STRIDE identified more fucosylated glycans (168 versus 78; 2.15-fold), sialylated glycans (240 versus 216; 1.11-fold), fucosylated and sialylated glycans (243 versus 111; 2.19-fold), and HexNAx + Hex compositions (86 versus 62; 1.39-fold) compared with DIA-NN (Fig. 5E). The mirror plot further showed that STRIDE added glycoforms across many proteins while retaining a shared core of glycoforms detected by both methods (Fig. 5D). Together, these results indicate that STRIDE improves across-run completeness for both acetyllysine and glycopeptide-enriched DIA datasets, with particularly strong gains in the reproducibility in the glycoform detection in human serum.

## Discussion

PTMs significantly expand protein functional diversity, but their identification and site-localization remain challenging due to the large combinatorial possibilities of modifiable amino acids.^2,44,45^ DIA analysis using unique ion signatures enable confident peptidoform identification.^14,19,46^ However, unique ion signatures may be low in abundance, and technical and biological heterogeneity across samples can compromise the quality of peptidoform peak-group assignment. STRIDE demonstrates significant improvements over DIA-NN and IPF for consistent identification and quantification of PTMs in DIA mass spectrometry. By aligning peak-groups across runs and propagating UIS signals, this method addresses a fundamental challenge in quantification of peptidoforms: the consistent assignment of peptidoforms to chromatographic peaks across multiple samples.

Performance on the benchmark dataset demonstrates the method’s sensitivity in site localization accuracy and correct peptidoform peak assignment. When considering both correct site localization and peak assignment, STRIDE reduced the false-negative rate from 68% with DIA-NN to 6% at a 5% false-positive rate. These improvements are particularly evident in the lower dilution runs, where signal quality deteriorates and DIA-NN either selects the wrong peak or fails to assign a peptidoform peak. STRIDE also improved the performance compared to IPF, reducing the false-negative rate from 14% to 6% when considering both correct site-localization and peak assignment. Propagating unique ion signature evidence across runs helps recover peptidoform assignments in runs with low-abundance signals.

Analyzing the phospho-enriched U2OS dataset demonstrates how STRIDE improves the analysis of biological experiments. By minimizing missing values and increasing signal consistency between replicates, STRIDE produces more complete quantification matrices. In the representative quantification matrix, missing values decreased from 38.6% with DIA-NN to 15.2% with STRIDE, while the mean number of quantified runs per phosphopeptide increased from 12.3 to 16.9. STRIDE also increased the number of highly reproducible phosphopeptides detected in at least 9 of 10 treated samples from 7,698 to 12,510, while reducing sporadic identifications detected in only 1-2 runs from 2,308 to 451. This boosts statistical power, allowing us to detect more differentially abundant phosphopeptides and enriched pathways than DIA-NN. Many of these signals matched the expected biology of nocodazole-induced spindle perturbation and checkpoint-maintained mitotic arrest. Shared DIA-NN and STRIDE differential phosphorylated proteins defined a core response across mitotic kinase, spindle motor and microtubule-organization modules, while STRIDE added phosphoprotein- and site-level coverage. Because these additional calls map to the same mitotic modules rather than separate responses, they support the interpretation that STRIDE expands the phosphosite-level view of the nocodazole-induced mitotic state.

The improvements for acetylated and glycosylated peptides further show that STRIDE offers advantages across different PTM types. In the acetyllysine dataset, STRIDE increased the number of complete acetyllysine peptide identifications across three replicate runs by 1.66-fold in the halfDIA condition and 1.71-fold in the fullDIA condition compared to DIA-NN. This was accompanied by a 44.6% reduction in missing quantification values in the halfDIA condition and a 51.9% reduction in the fullDIA condition. In the human serum glycopeptide dataset, STRIDE increased site-specific glycan identifications from 467 to 737 and increased the number of detected in all three replicated form 8 to 460. STRIDE also reduced the missing quantification values by 63.1% at the glycopeptide level, 63.0% at the site-specific glycan level, and 81.5% at the protein glycosite level. These results indicate that the method improves reproducibility and quantification completeness, especially when the same modified species are expected to be quantified consistently across replicate runs.

Overall, our results demonstrate that STRIDE improves peptidoform proteomics analysis in the following ways: increased identification sensitivity, reduced false-negative rates, improved site-localized peak assignment, and more consistent quantification. The method may be particularly valuable in complex, multi-run experimental designs, where alignment and signal propagation can improve quantification of low-abundance species and increase statistical power for differential analyses.

## Methods

### Raw Spectral File Conversion

Vendor spectral files were converted to the open mzML format using MSConvert (dockerhub: chambm/pwiz-skyline-i-agree-to-the-vendor-licenses:1493f57e824b) with centroiding, remaining settings were left as defaults. All files were converted this way unless stated otherwise. The synthetic phospho dataset files were retrieved from PRIDE archive with the accession PXD004573, the U2OS phospho-enriched dataset files were retrieved from PRIDE with accession PXD017476, the mouse liver acetylated-enriched dataset files were retrieved from MassIVE with the accession MSV000080189, and the human serum glycosylated-enriched dataset was retrieved from ProteomeXchange with the accession PXD023980.

### Spectral Library Generation

Data-independent acquisition (DIA) files were processed with DIA-Umpire to generate pseudo-DDA spectra, using the defaults with the *Mass Defect Filter* turned off for PTMs. The resulting pseudo-DDA spectra and, if available, data-dependent acquisition data (DDA) were searched using MSFragger using the reasonable default settings, with the added variable modifications specific for the dataset being processed (i.e. STY for phosphorylation, nK for acetylation). Resulting identified peptide spectrum matches (PSMs) were statistically validated downstream using peptide prophet, PTM prophet and protein prophet. Subsequently, confident PSMs were used to create a spectral transition library using easyPQP. These steps were all performed using the FragPipe software. The final peptide query parameter (PQP) library was generated using OpenSwathAssayGenerator and OpenSwathDecoyGenerator to append theoretical identifying transitions and alternative peptidoforms, as described in Rosenberger, G. *et al.*^14^ For retention time calibration, linear and non-linear PQPs were generated using the run-specific PSMs identified in the pseudo-DDA spectra and processed using easyPQP. Each dataset used the above mentioned approach for spectral library generation, with the exception of the human serum glycosylated-enriched dataset. For this dataset, we used the published semi-empirical library as described in Yang, Y. *et al.*^43^ as the library was already in the desired peptide query parameter format required by OpenSwathWorkflow.

### Peak-Group Feature Identification

OpenSwathWorkflow was used to identify transition-group chromatographic peak features in the DIA spectra using the PQP library generated in the previous step. The following parameters were commonly set across the datasets:

- enable_ms1 -enable_ipf
- min_upper_edge_dist 1
- RTNormalization:alignmentMethod lowess
- RTNormalization:estimateBestPeptides
- Scoring:Scores:use_uis_scores.

The following parameters below were set specifically for each dataset.

Synthetic phospho dataset:

- mz_extraction_window 30
- mz_extraction_window_ms1 20
- mz_correction_function regression_delta_ppm
- rt_extraction_window 300
- RTNormalization:lowess:span 0.1
- RTNormalization:outlierMethod none
- Scoring:TransitionGroupPicker:background_subtraction exact

U2OS phospho-enriched dataset:

- mz_extraction_window 50
- mz_extraction_window_ms1 50
- mz_correction_function regression_delta_ppm
- extra_rt_extraction_window 100
- RTNormalization:MinBinsFilled 5
- RTNormalization:lowess:span 0.1
- RTNormalization:outlierMethod none
- Scoring:TransitionGroupPicker:background_subtraction exact

Mouse liver acetylated-enriched dataset:

- mz_extraction_window 30
- mz_extraction_window_ms1 20
- mz_correction_function regression_delta_ppm
- rt_extraction_window 300
- RTNormalization:lowess:span 0.1
- RTNormalization:outlierMethod none
- Scoring:TransitionGroupPicker:background_subtraction exact

Human serum glycosylated-enriched dataset:

- mz_extraction_window 20
- mz_extraction_window_ms1 10
- mz_correction_function quadratic_regression_delta_ppm
- RTNormalization:NrRTBins 6
- RTNormalization:MinBinsFilled 4
- RTNormalization:lowess:span 0.01

### Statistical Validation of Peak-Group Features and Inference of Peptidoforms

PyProphet (v2.3.0) was used for semi-supervised learning and error-rate estimation of the features identified by OpenSWATH, on the peak-group (ms1ms2) and individual transition level. For aligned results, scoring was applied to the aligned features as well. For inference of peptidoforms, ms1 and ms2 precursor data was disabled, only allowing for transition level evidence to guide the inference. For aligned results, the cross-run signal propagation was enabled to propagate a maximum of transition level signal (posterior error probability) of 0.5. Subsequently, global peptide-level inference was performed using the default parameters. For the N-glycosylation dataset, similar steps were performed, except using the scoring and glycoform inference described in Yang, Y. *et al.*^43^ with the added alignment and signal propagation method.

### DIA-NN Analysis

All analyses with DIA-NN used version DIA-NN v2.1, which supports peptidoform assessment. The recommended default parameters were used as suggested in DIA-NN’s documentation:

--reannotate --cut K*,R* --missed-cleavages 1 --unimod4 --var-mods 3 --var-mod UniMod:35,15.994915,M --var-mod UniMod:1,42.010565,*n --var-mod UniMod:21,79.966331,STY --use-quant --peptidoforms --reanalyse --rt-profiling --report-decoys

## Data Availability

The synthetic phospho dataset files were retrieved from PRIDE archive with the accession PXD004573, the U2OS phospho-enriched dataset files were retrieved from PRIDE with accession PXD017476, the mouse liver acetylated-enriched dataset files were retrieved from MassIVE with the accession MSV000080189, and the human serum glycosylated-enriched dataset was retrieved from ProteomeXchange with the accession PXD023980. The ARYCAL software for chromatogram alignment is open source and freely available under a BSD-3 clause license on GitHub (https://github.com/singjc/arycal), and available with Python bindings via the PyPi repository. Targeted data extraction performed with OpenSWATH (OpenMS >v3.4) is open source and freely available under a BSD-3 clause license on GitHub (https://github.com/OpenMS/OpenMS). Statistical scoring and peptidoform inference performed with PyProphet (>2.3) is open source and freely available under a BSD-3 clause license on GitHub (https://github.com/PyProphet/pyprophet).

## Contributions

J.C.S. and H.L.R. designed the study. J.C.S. designed and developed the method. H.L.R. supervised the project. S.G. provided advice on the alignments of extracted ion chromatograms. All authors contributed to the writing of the manuscript. All authors read and approved the final manuscript.

## Acknowledgments

J.C.S. was supported by the European Research Area Network Personalized Medicine Cofund (PerProGlio, #506078) and CIHR (#507496).

