## Supplemental Information for "STRIDE: Signal Transfer and Aligned Ion-Peak Discrimination for Peptidoform Evidence for Consistent Quantification of Site-Localized Post-Translational Modifications in Large-Scale DIA-MS"

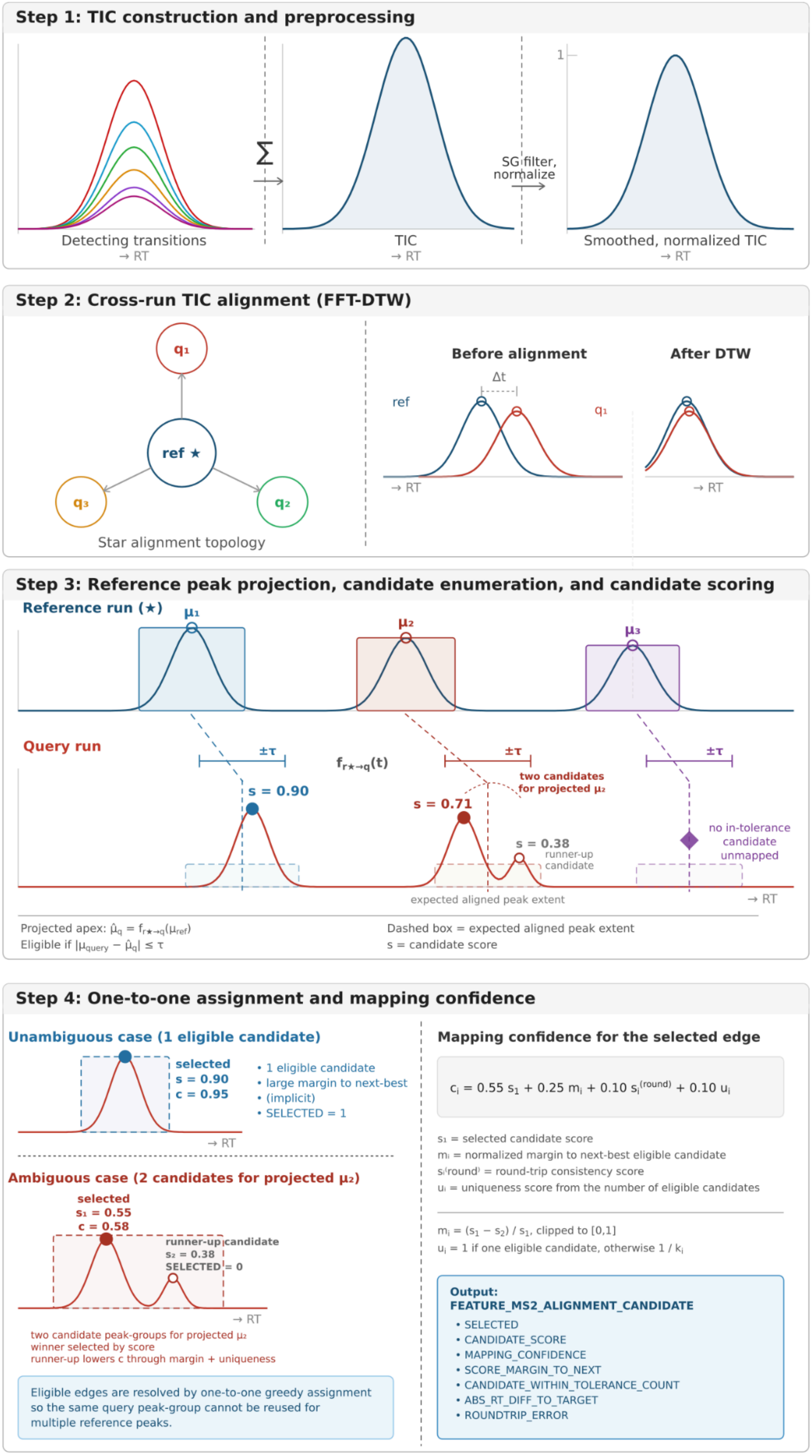

Step 1: TIC construction and preprocessing
Detecting transitions
→ RT
Σ
TIC
→ RT
SG filter, normalize
1
Smoothed, normalized TIC
→ RT
Step 2: Cross-run TIC alignment (FFT-DTW)
$q_1$
ref ★
$q_3$
$q_2$
Star alignment topology
Before alignment
After DTW
Δt
ref
$q_1$
→ RT
→ RT
Step 3: Reference peak projection, candidate enumeration, and candidate scoring
Reference run (★)
$\mu_1$
$\mu_2$
$\mu_3$
Query run
±τ
±τ
±τ
$f_{r\star \to q}(t)$
two candidates for projected $\mu_2$
s = 0.90
s = 0.71
s = 0.38
runner-up candidate
no in-tolerance candidate unmapped
expected aligned peak extent
→ RT
Projected apex: $\hat{\mu}_q = f_{r\star \to q}(\mu_{ref})$
Eligible if $|\mu_{query} - \hat{\mu}_q| \leq \tau$
Dashed box = expected aligned peak extent
s = candidate score
Step 4: One-to-one assignment and mapping confidence
Unambiguous case (1 eligible candidate)
selected
s = 0.90
c = 0.95
• 1 eligible candidate
• large margin to next-best
• (implicit)
• SELECTED = 1
→ RT
Ambiguous case (2 candidates for projected $\mu_2$)
selected
$s_1$ = 0.55
c = 0.58
runner-up candidate
$s_2$ = 0.38
SELECTED = 0
→ RT
two candidate peak-groups for projected $\mu_2$
winner selected by score
runner-up lowers c through margin + uniqueness
Eligible edges are resolved by one-to-one greedy assignment so the same query peak-group cannot be reused for multiple reference peaks.
Mapping confidence for the selected edge
$c_i = 0.55\, s_1 + 0.25\, m_i + 0.10\, s_i^{(round)} + 0.10\, u_i$
$s_1$ = selected candidate score
$m_i$ = normalized margin to next-best eligible candidate
$s_i^{(round)}$ = round-trip consistency score
$u_i$ = uniqueness score from the number of eligible candidates
$m_i = (s_1 - s_2) / s_1$, clipped to [0,1]
$u_i$ = 1 if one eligible candidate, otherwise 1 / $k_i$
Output:
FEATURE_MS2_ALIGNMENT_CANDIDATE
• SELECTED
• CANDIDATE_SCORE
• MAPPING_CONFIDENCE
• SCORE_MARGIN_TO_NEXT
• CANDIDATE_WITHIN_TOLERANCE_COUNT
• ABS_RT_DIFF_TO_TARGET
• ROUNDTRIP_ERROR

**Supplementary Figure 1. Overview of the across run dynamic chromatogram alignment (ARYCAL) and confidence-aware peak-group mapping workflow.**
For each precursor, ARYCAL first extracts detecting-transition chromatograms in each run and collapses them into a total ion chromatogram (TIC), followed by Savitzky-Golay smoothing and intensity normalization **(Step 1)**. Smoothed run-level TICs are then aligned to a common reference run using a star-alignment strategy with FFT-initialized dynamic time warping (FFT-DTW), yielding a nonlinear retention-time mapping between the reference and each query run **(Step 2)**. Reference-run peak-groups are projected into each query run using the learned RT mapping, producing a projected apex $\hat{\mu}_q$ and an expected aligned peak extent; all same-precursor query peak-groups are enumerated and scored as candidate mappings, and candidates are considered eligible when their apex RT satisfies $|\mu_{\text{query}} - \hat{\mu}_q| \leq \tau$ **(Step 3)**. Candidate scores are computed from RT agreement, projected-versus-observed boundary overlap, peak-width similarity, and optional peak-group metadata (e.g. rank and q-value). Eligible candidate edges are then resolved by one-to-one greedy assignment so that the same query peak-group cannot be reused for multiple reference peaks. For each selected edge, a final mapping confidence is computed as a weighted combination of the selected candidate score, the normalized score margin to the next-best eligible candidate, round-trip RT consistency, and a uniqueness term reflecting the number of in-tolerance candidates **(Step 4)**. The resulting candidate-level output table, FEATURE_MS2_ALIGNMENT_CANDIDATE, stores all evaluated reference-query peak-group pairs together with the selected mapping and mapping-confidence metrics for downstream filtering during across-run signal propagation.

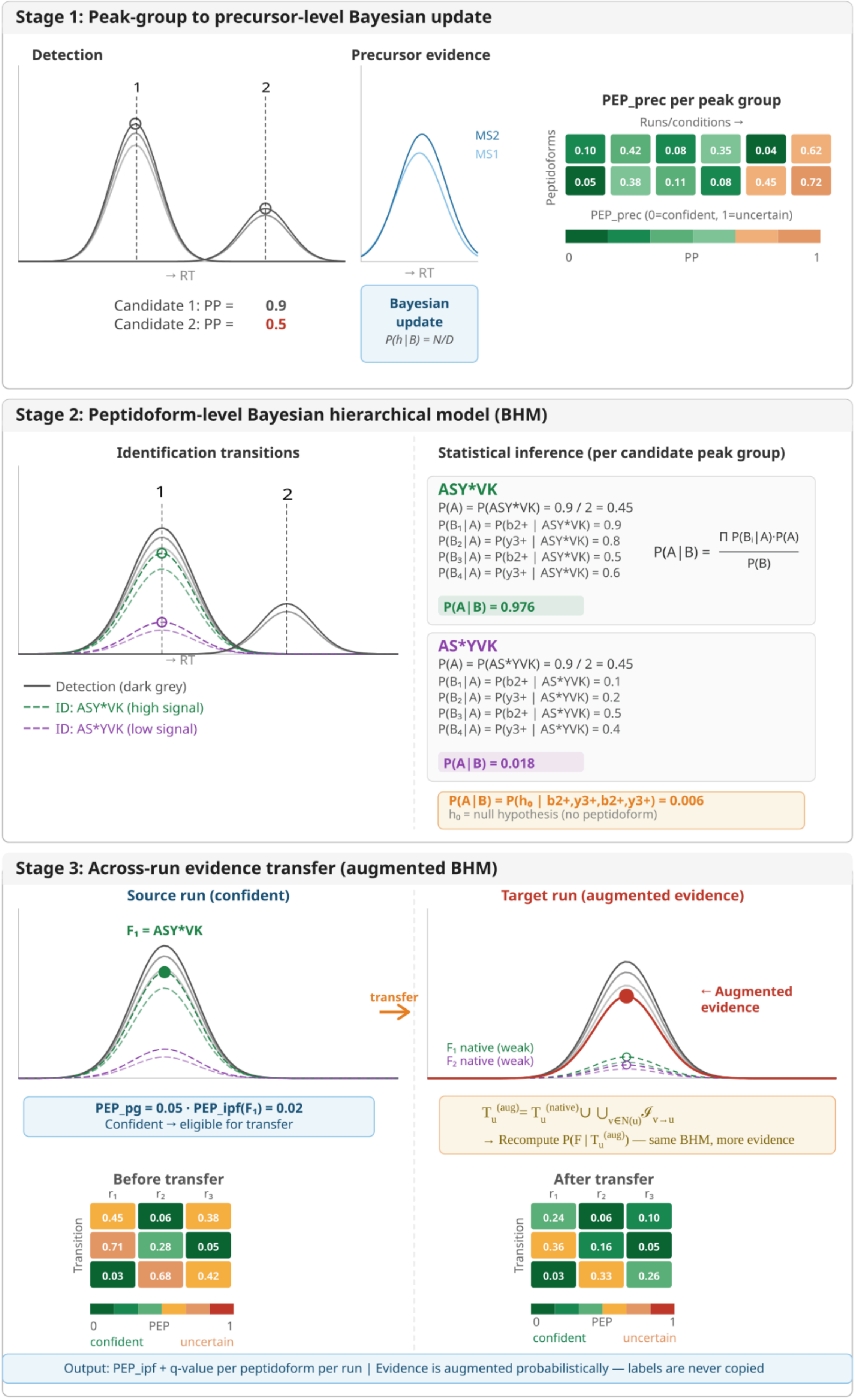

Stage 1: Peak-group to precursor-level Bayesian update
Detection
Precursor evidence
1
2
→ RT
Candidate 1: PP = 0.9
Candidate 2: PP = 0.5
MS2
MS1
→ RT
Bayesian update
P(h | B) = N/D
PEP_prec per peak group
Runs/conditions →
Peptidoforms
0.10 0.42 0.08 0.35 0.04 0.62
0.05 0.38 0.11 0.08 0.45 0.72
PEP_prec (0=confident, 1=uncertain)
0 PP 1
Stage 2: Peptidoform-level Bayesian hierarchical model (BHM)
Identification transitions
Statistical inference (per candidate peak group)
→ RT
Detection (dark grey)
ID: ASY*VK (high signal)
ID: AS*YVK (low signal)
ASY*VK
P(A) = P(ASY*VK) = 0.9 / 2 = 0.45
P(B1 | A) = P(b2+ | ASY*VK) = 0.9
P(B2 | A) = P(y3+ | ASY*VK) = 0.8
P(B3 | A) = P(b2+ | ASY*VK) = 0.5
P(B4 | A) = P(y3+ | ASY*VK) = 0.6
P(A | B) = Π P(Bi | A)·P(A) / P(B)
P(A | B) = 0.976
AS*YVK
P(A) = P(AS*YVK) = 0.9 / 2 = 0.45
P(B1 | A) = P(b2+ | AS*YVK) = 0.1
P(B2 | A) = P(y3+ | AS*YVK) = 0.2
P(B3 | A) = P(b2+ | AS*YVK) = 0.5
P(B4 | A) = P(y3+ | AS*YVK) = 0.4
P(A | B) = 0.018
P(A | B) = P(h0 | b2+,y3+,b2+,y3+) = 0.006
h0 = null hypothesis (no peptidoform)
Stage 3: Across-run evidence transfer (augmented BHM)
Source run (confident)
Target run (augmented evidence)
F1 = ASY*VK
transfer
← Augmented evidence
F1 native (weak)
F2 native (weak)
PEP_pg = 0.05 · PEP_ipf(F1) = 0.02
Confident → eligible for transfer
Tu(aug) = Tu(native) ∪ ∪v∈N(u) 𝒯v→u
→ Recompute P(F | Tu(aug)) — same BHM, more evidence
Before transfer
After transfer
r1 r2 r3
Transition
0.45 0.06 0.38
0.71 0.28 0.05
0.03 0.68 0.42
0.24 0.06 0.10
0.36 0.16 0.05
0.03 0.33 0.26
0 PEP 1
confident uncertain
Output: PEP_ipf + q-value per peptidoform per run | Evidence is augmented probabilistically — labels are never copied

**Supplementary Figure 2. Overview of the IPF signal propagation framework for across-run peptidoform inference.** The IPF workflow proceeds in three stages. In **Stage 1**, detection-level precursor evidence from MS1 and MS2 peak-group signals is combined within each run and candidate peak-group using a Bayesian update to produce a precursor-level posterior error probability ($\mathrm{PEP}_{\mathrm{prec}}$), which summarizes how confidently a given peak-group supports the presence of the precursor. In **Stage 2**, identification transitions are evaluated within each candidate peak-group using a Bayesian hierarchical model (BHM) to infer peptidoform identity from the observed fragment-ion evidence. This yields posterior probabilities and peptidoform-specific error estimates ($\mathrm{PEP}_{\mathrm{ipf}}$) for each candidate peak-group, while retaining an explicit null hypothesis when no peptidoform is sufficiently supported. In **Stage 3**, across-run signal propagation augments the evidence available in weak target runs using aligned, high-confidence peptidoform evidence from source runs. Rather than copying discrete labels across runs, the method transfers probabilistic evidence into the target run and recomputes the same hierarchical Bayesian model using the augmented evidence set. As a result, native weak signals can be reinforced by concordant across-run evidence, leading to improved peptidoform confidence and downstream q-values, while preserving uncertainty and allowing unsupported hypotheses to remain unassigned.

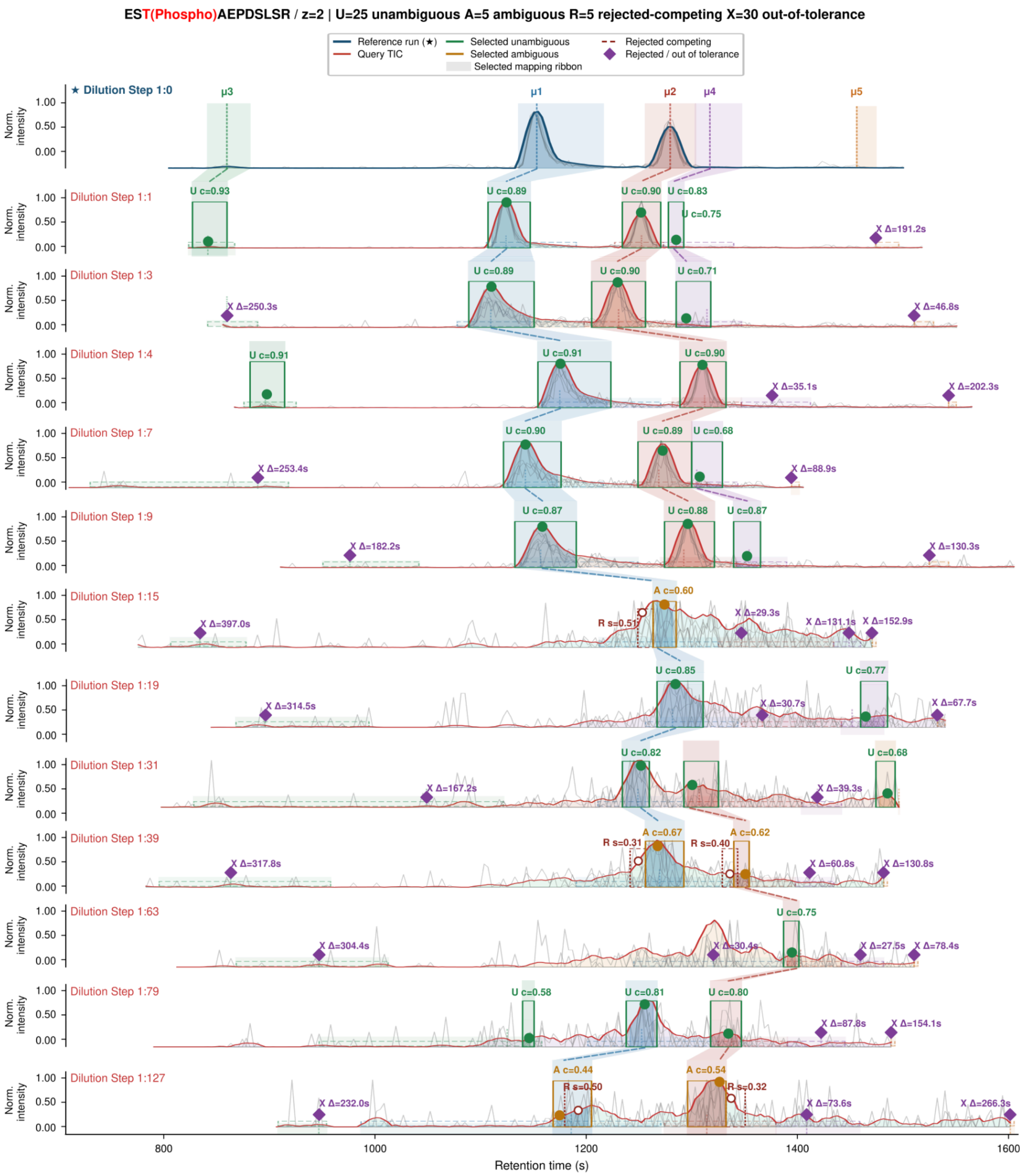


**Supplementary Figure 3. Peptidoform visualization of unambiguous, ambiguous, rejected-competing, and out-of-tolerance alignment candidates across runs.** Shown is the ARYCAL candidate-level mapping output for the precursor EST(Phospho)AEPDSLSR, charge state 2, across a dilution-series experiment. The top panel is the reference run, in which reference peak-groups ($\mu_1 \cdots \mu_5$) are defined. In each query run, the red trace shows the run-level smoothed TIC and grey traces show the underlying detecting-transition chromatograms. Reference peak-groups are projected into each query run using the learned RT mapping, yielding a projected target RT and an expected aligned peak extent. Green solid windows indicate selected unambiguous mappings, for which only one in-tolerance candidate was available. Orange solid windows indicate selected ambiguous mappings, in which multiple in-tolerance candidates were present and the selected edge won by candidate score but retained lower confidence due to reduced margin and uniqueness. Red dashed windows mark rejected competing candidates that were within tolerance but lost to a higher-scoring selected candidate. Purple diamonds indicate out-of-tolerance cases, in which no eligible in-tolerance candidate was available for that projected reference peak. Translucent ribbons connect selected mappings across adjacent runs, illustrating the across-run correspondence used for downstream signal propagation. The title summary reports the total number of selected unambiguous mappings (U), selected ambiguous mappings (A), rejected competing candidates (R), and out-of-tolerance cases (X) for this precursor across all displayed runs.

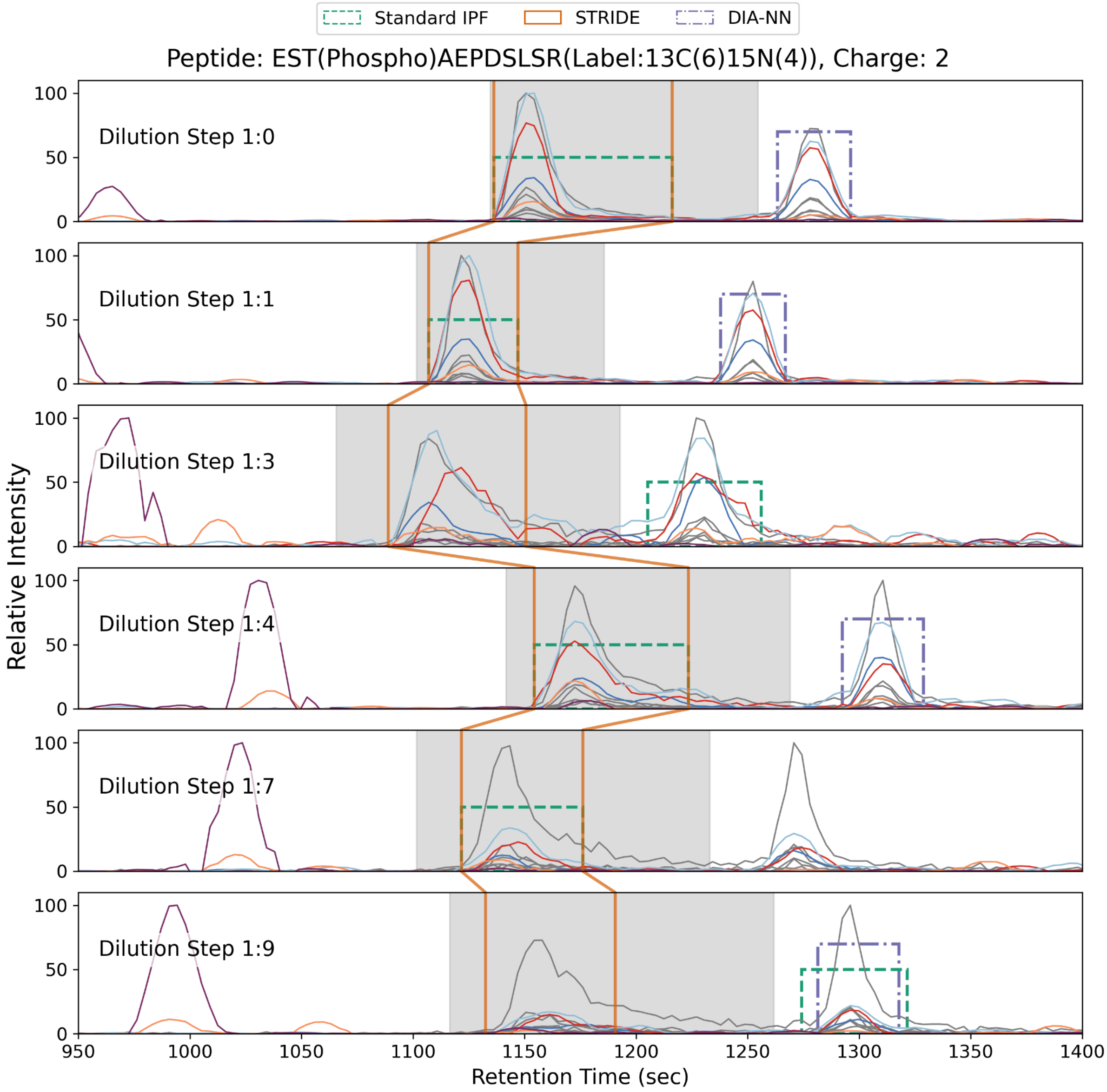


**Supplementary Figure 4. Consistent Peptidoform-Peak Assignment.** Example extracted ion chromatograms for phosphorylated peptide EST(Phospho)AEPDSLSR across the first 6 dilution steps. The last 7 dilution step runs did not have any peaks detected by either standard IPF or STRIDE. The grey traces represent the empirical peak-group detecting transitions, and the colored traces represent unique-ion signatures for the b3 and y9 ions. The grey shaded region represents the manual annotated region from Skyline, the green dashed rectangles represent the peak boundaries assigned by the standard inference of peptidoform (IPF) workflow at 5% false discovery rate (FDR), and the orange boundaries connected across the runs represent peak boundaries assigned by STRIDE at 5% FDR. The purple rectangle boundaries represent peptidoform peak identification by DIA-NN at 5% FDR (1 - site localization probability).

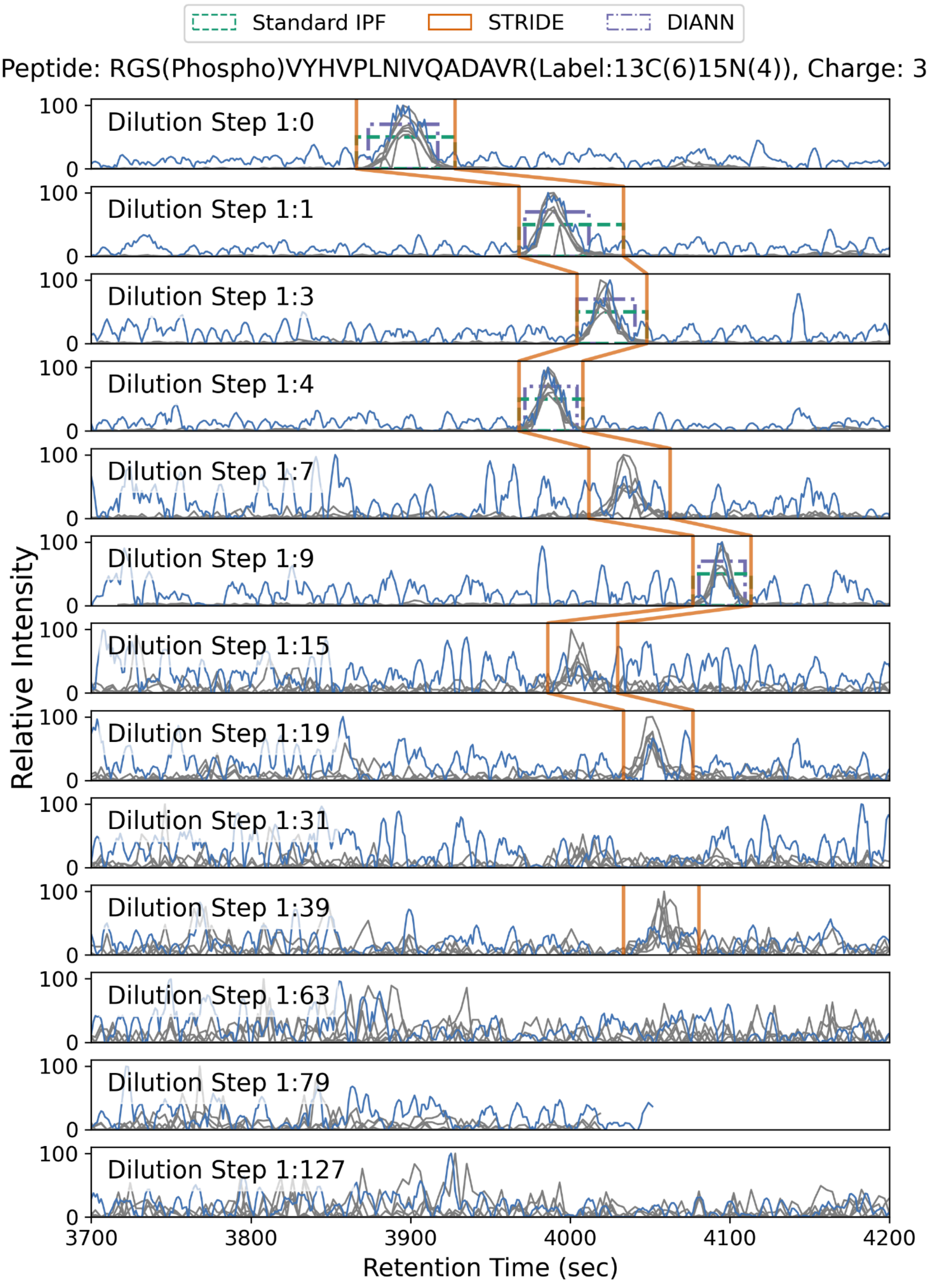


**Supplementary Figure 5. Recovery of missed peaks in lower dilutions.** Example of extracted ion chromatograms for phosphorylated peptide

RGS(Phospho)VYHPLNIVQADAVR across all 13 dilutions. The grey traces represent the empirical peak-group detecting transitions, and the blue trace represents the unique-ion signatures for the b3 ions. The green dashed rectangles represent the peak boundaries assigned by the standard inference of peptidoform (IPF) workflow at 5% false discovery rate (FDR), and the orange boundaries connected across the runs represent peak boundaries assigned by STRIDE at 5% FDR. The purple rectangle boundaries represent peptidoform peak identification by DIA-NN at 5% FDR.

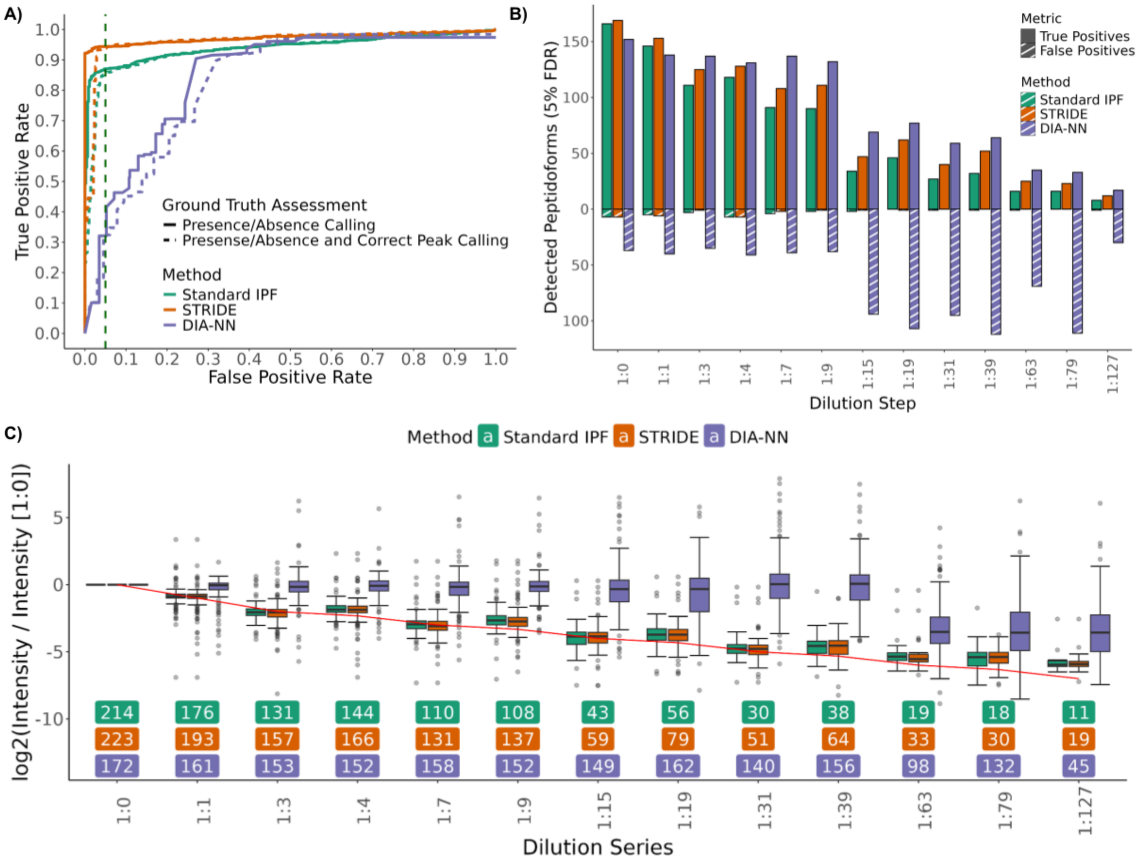


**Supplementary Figure 6. Comparison to DIA-NN on Benchmark Synthetic Phosphopeptide Dataset. A)** The receiver operating characteristics (ROC) curve shows the identification performance of the standard IPF workflow, STRIDE and DIA-NN. The solid line shows the performance when assessing only for the correct peptidoform identification, standard IPF (green) at 5% false positive rate (FPR) has 87% true positive rate (TPR), STRIDE (orange) at 5% FPR has 94% TPR and DIA-NN (purple) at 5% FPR has 32% TPR.  The dashed lines show the performance of correctly identifying the right peptidoform as well as correctly assigning it to the correct peak (from manual annotations). Standard IPF at 5% FPR achieves 86% TPR, STRIDE at 5% FPR achieves 94 % TPR, and DIA-NN at 5% FPR achieves 32% TPR. **B)** The bar plot shows the number of identifications per dilution step at a 5% FDR controlled for the peptidoform level. The green bars represent the standard IPF workflow,  the orange bars represent STRIDE, and the purple bars represent DIA-NN. The solid bars represent the true positives, and the striped bars represent the false positives. **C)** Quantification of the peptidoform assigned peak-groups at 5% site-localization FDR, normalized to the highest concentration run (1:0). The red line indicates the ground truth of the dilution series. The green boxplots represent quantification from the standard IPF workflow, the orange box plots represent quantification from STRIDE, and the purple box plots represent quantification from DIA-NN. The numbers below the boxplots represent the number of phosphopeptides identified by each method, color code matches the corresponding methods boxplot color.

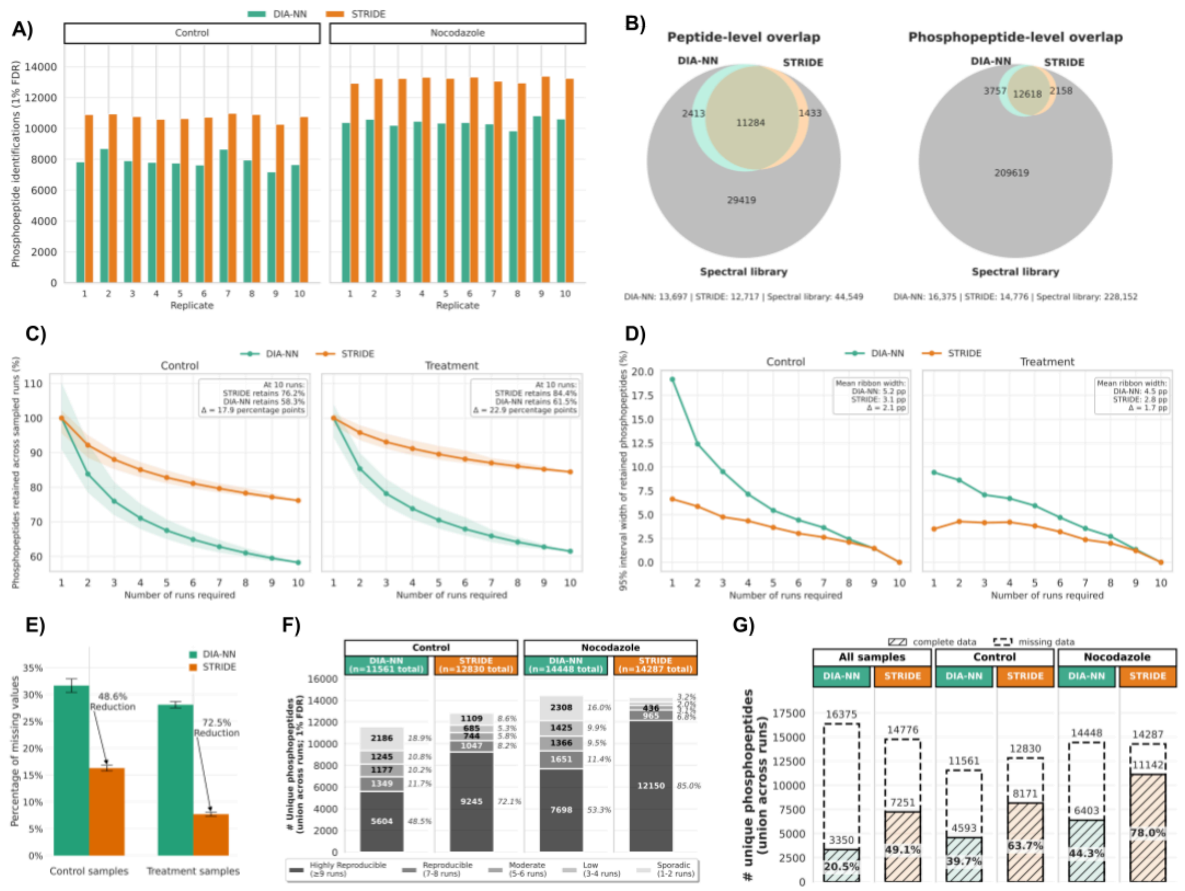


**Supplementary Figure 7. STRIDE improves phosphopeptide identification completeness and run-to-run consistency in the U2OS nocodazole experiment.** **A)** Number of phosphopeptide identifications per replicate for DIA-NN and STRIDE, separated by control and nocodazole-treated samples. **B)** Venn diagrams showing peptide-level and phosphopeptide-level overlap between DIA-NN, STRIDE, and the spectral library. **C)** Identification consistency across random subsets of runs. The y-axis shows the percentage of phosphopeptides retained as the number of required runs increases; shaded regions show the 95% interval across repeated random sampling. **D)** Width of the 95% interval from the run-sampling analysis in panel C, shown as a measure of run-to-run variability in identification consistency. **E)** Percentage of missing phosphopeptide intensity values for DIA-NN and STRIDE in control and nocodazole-treated samples. Error bars indicate variation across samples. **F)** Reproducibility of phosphopeptide identifications across runs, grouped by condition and method. Stacked bars show the number of unique phosphopeptides detected in the indicated number of runs. **G)** Complete and incomplete phosphopeptide quantification across all samples, control samples, and nocodazole-treated samples. Hatched bars indicate phosphopeptides detected in all runs, and dashed outlines indicate phosphopeptides detected in at least one run. Green indicates DIA-NN and orange indicates STRIDE.

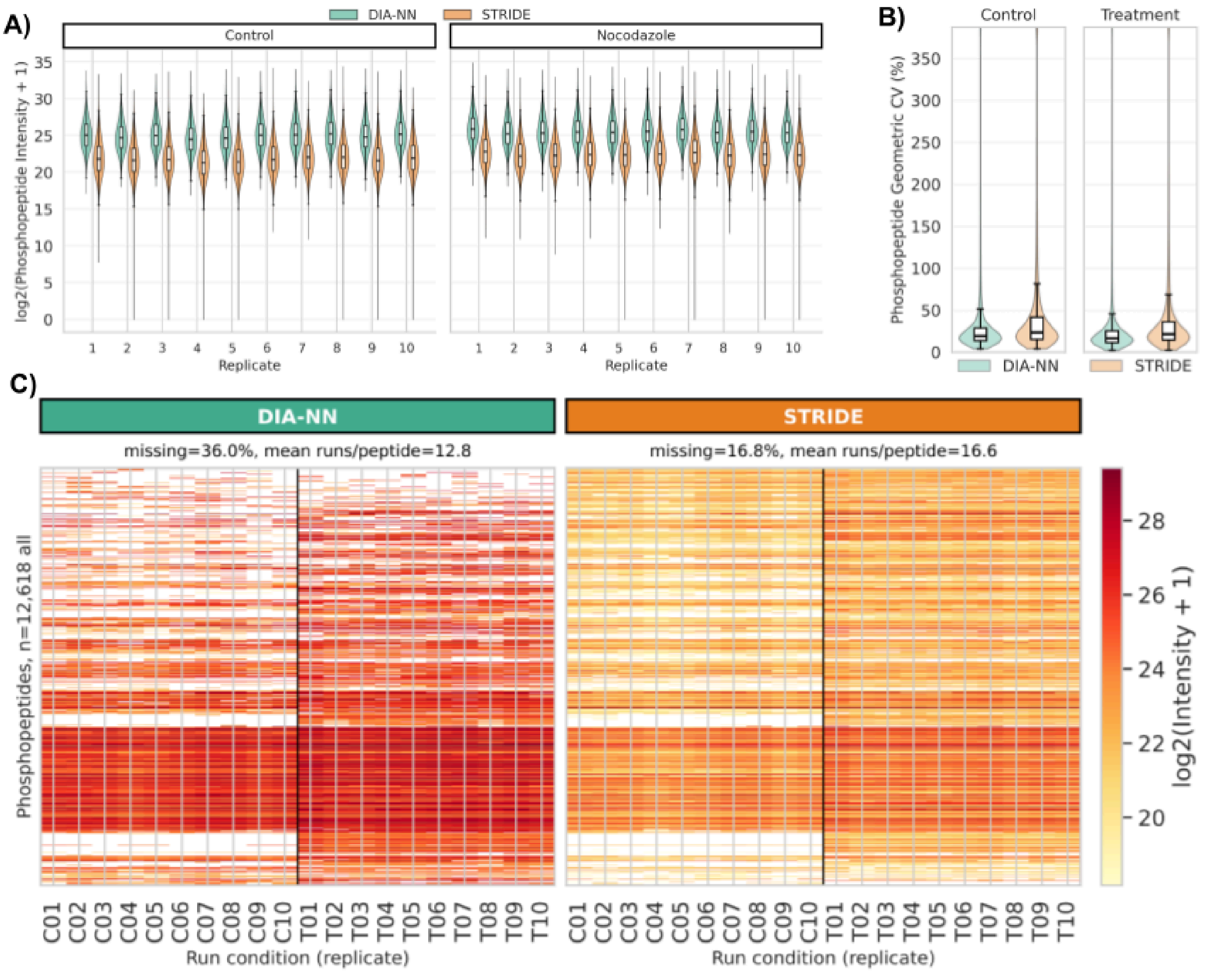


**Supplementary Figure 8. Phosphopeptide intensity distributions and quantification matrices in the U2OS nocodazole experiment.**
**A)** Distribution of log2-transformed phosphopeptide intensities for DIA-NN and STRIDE across individual replicates, separated by control and nocodazole-treated samples. Violin plots show the intensity distribution for each replicate, with inner boxplots indicating the median and interquartile range. **B)** Distribution of phosphopeptide geometric coefficients of variation (CV) for DIA-NN and STRIDE in control and nocodazole-treated samples. **C)** Phosphopeptide-by-run quantification matrices for all intersecting phosphopeptides for DIA-NN and STRIDE. Rows represent phosphopeptides and columns represent samples ordered by condition and replicate. Color indicates log2-transformed intensity, and white entries indicate missing values. The text above each matrix reports the percentage of missing values and the mean number of runs in which each phosphopeptide was quantified.

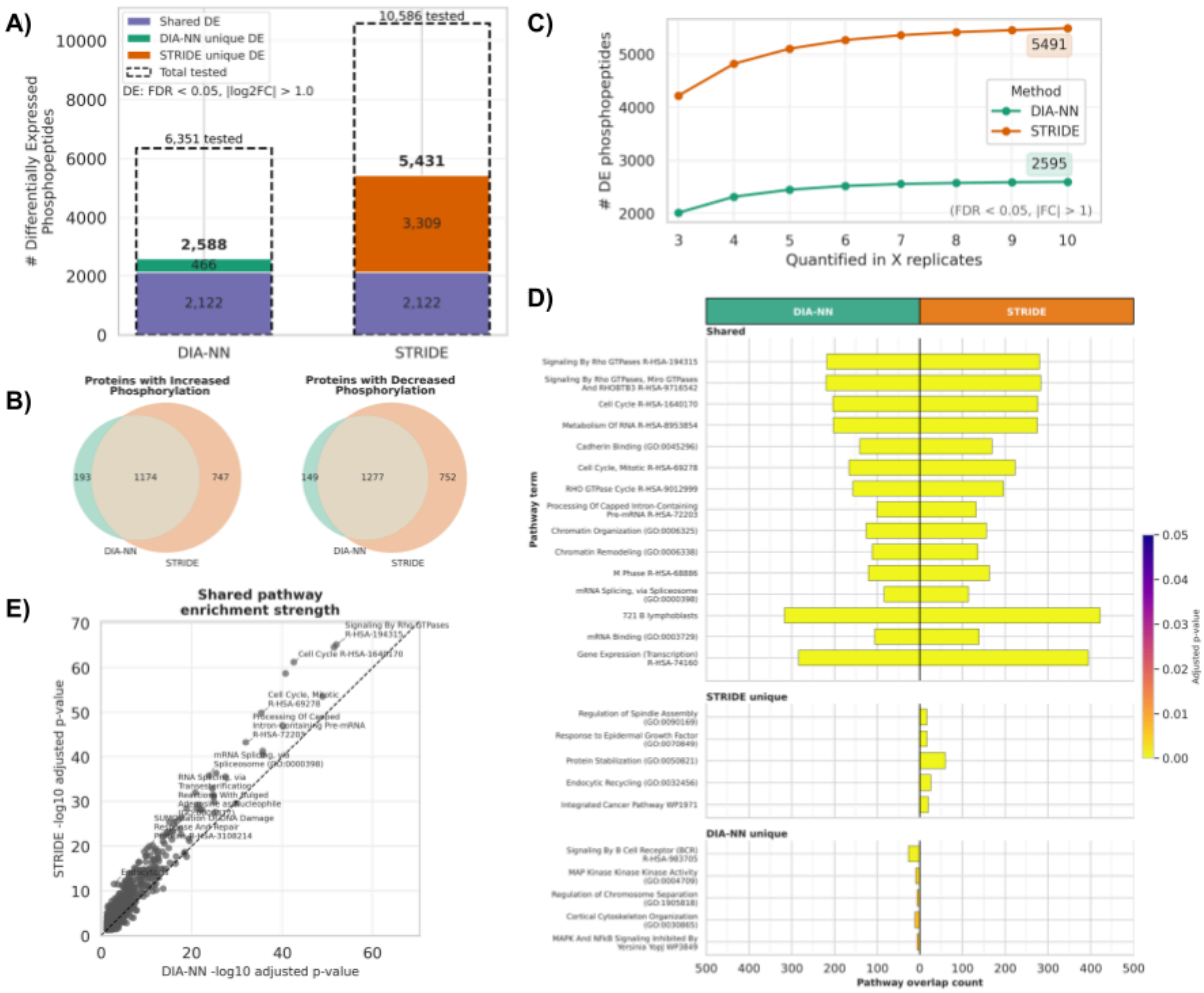


**Supplementary Figure 9. Increasing Number of Differentially Expressed Phosphopeptides and Pathway Enrichment. A)** Differentially expressed phosphopeptides identified by DIA-NN and STRIDE using an FDR threshold of 0.05 and an absolute log2 fold-change threshold greater than 1. Stacked bars show phosphopeptides shared between methods, unique to DIA-NN, and unique to STRIDE; dashed outlines indicate the total number of phosphopeptides tested by each method. **B)** Overlap of proteins with increased or decreased phosphorylation, based on differentially expressed phosphopeptides identified by DIA-NN and STRIDE. **C)** Number of differentially expressed phosphopeptides retained after requiring quantification in at least the indicated number of replicates. **D)** Mirrored pathway enrichment plot comparing DIA-NN and STRIDE. Shared pathways are shown in the top panel, STRIDE-unique pathways in the middle panel, and DIA-NN-unique pathways in the bottom panel. Bar length indicates pathway overlap count, and color indicates adjusted p-value. **E)** Shared pathway enrichment strength comparison between DIA-NN and STRIDE, shown as -log10 adjusted p-values for pathways detected by both methods. The dashed line indicates equal enrichment strength between methods.

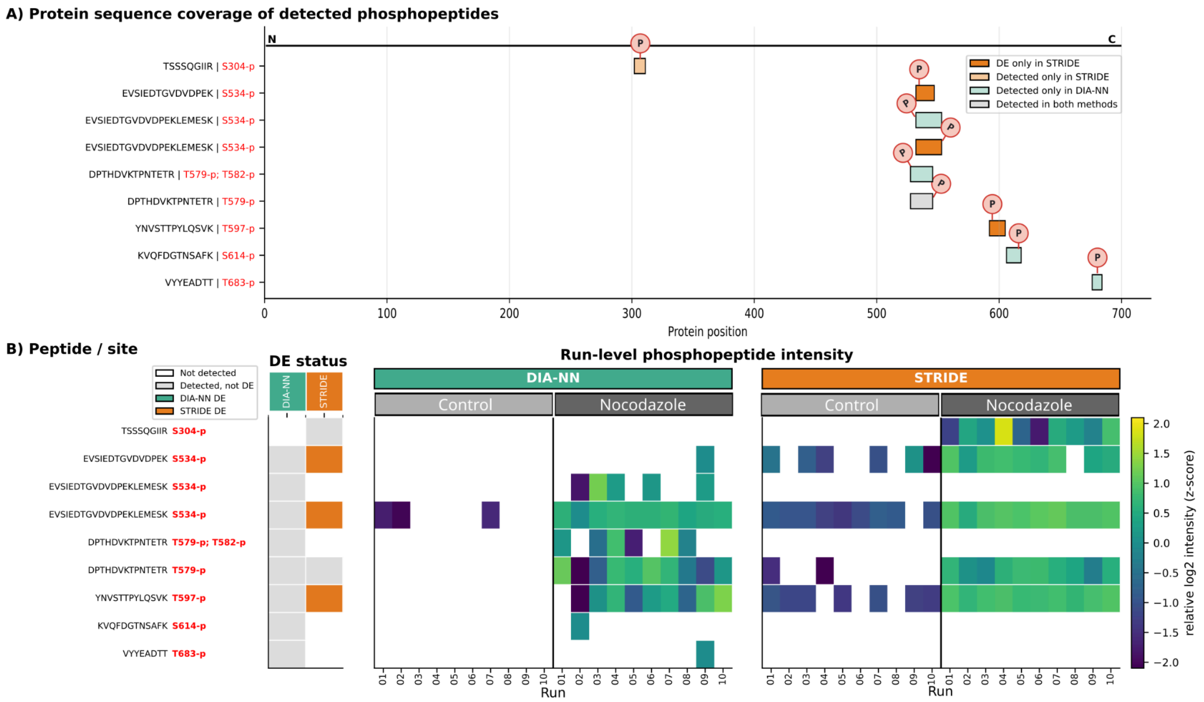


**Supplementary Figure 10. STRIDE reveals a coordinated multi-site CKAP2 phosphorylation pattern in the U2OS nocodazole experiment. A)** Protein sequence coverage of detected CKAP2 phosphopeptides. The horizontal line represents CKAP2 from the N- to C-terminus, rows show detected phosphopeptide/site assignments, and boxes indicate the corresponding protein positions. Orange indicates differentially expressed phosphopeptides detected only by STRIDE, tan indicates phosphopeptides detected only by STRIDE but not differentially expressed, teal indicates phosphopeptides detected only by DIA-NN, and grey indicates phosphopeptides detected by both methods. Phosphorylation sites are marked with *P*. **B)** Method-level differential-expression status and run-level relative phosphopeptide intensities for DIA-NN and STRIDE. Rows correspond to the phosphopeptide/site assignments in panel A, and columns represent individual control and nocodazole-treated runs. Intensities are shown as row-wise z-scores of log2 phosphopeptide intensity; white cells indicate missing values. The left annotation indicates whether each phosphopeptide was not detected, detected but not differentially expressed, differentially expressed by DIA-NN, or differentially expressed by STRIDE. Several CKAP2 phosphopeptides show increased abundance in nocodazole-treated samples, with more complete run-level coverage by STRIDE.

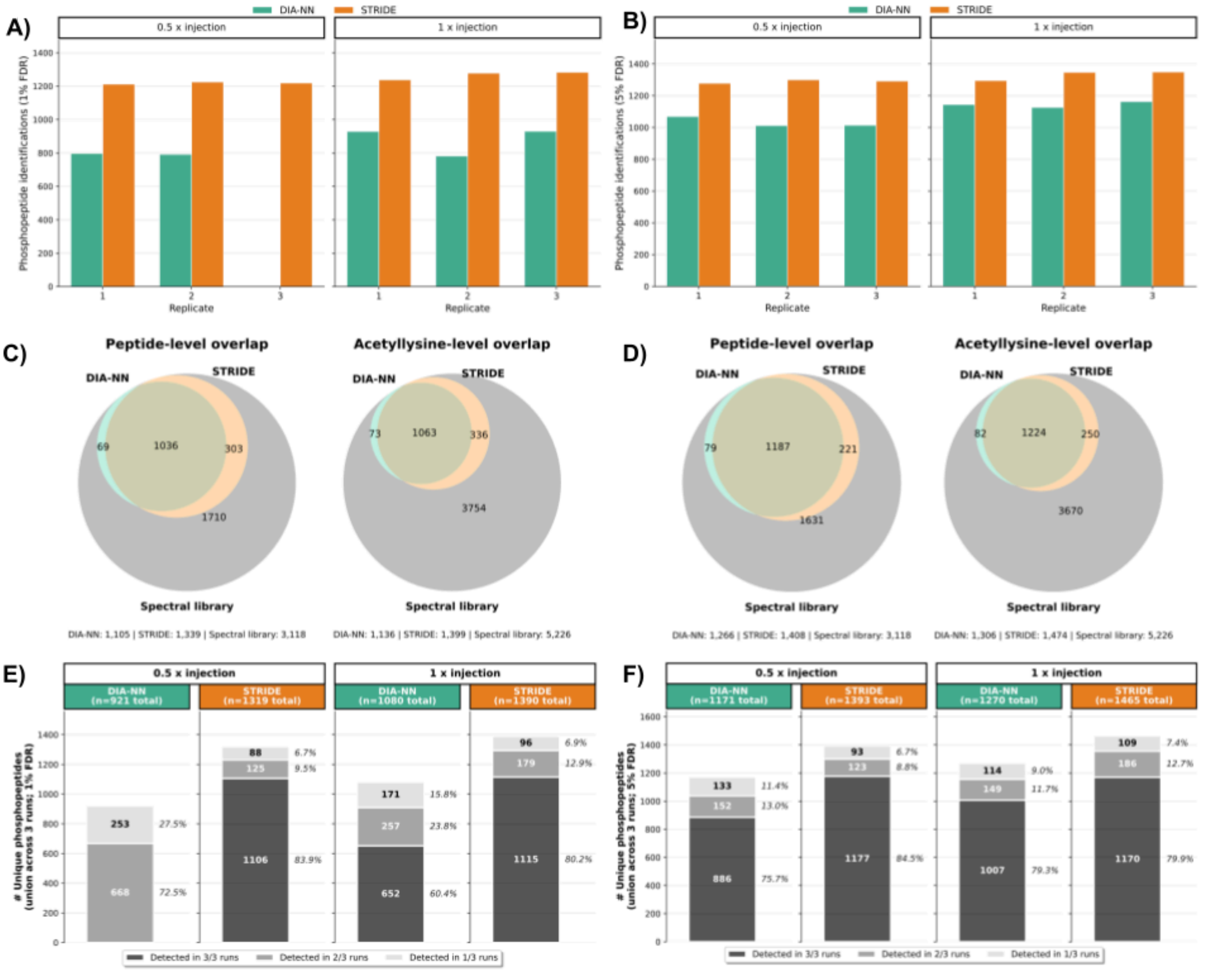


**Supplementary Figure 11. STRIDE increases acetyllysine identification coverage and reproducibility. A-B)** Number of acetyllysine peptide identifications per replicate for DIA-NN and STRIDE at 1% site-localization FDR (**A**) and 5% site-localization FDR (**B**). Results are shown separate for the 0.5 x injection and 1 x injection conditions. **C-D)** Overlap between DIA-NN, STRIDE and the spectral library at the peptide and acetyllysine-site levels at 1% FDR (**C)** and 5% FDR (**D**). **E-F)** Reproducibility of acetyllysine identifications across replicate runs at 1% FDR (**E**) and 5% FDR (**F**). Stacked bars show the number of unique acetyllysine peptides detected in 3/3/, 2/2 and 1/3 replicate runs for each injection condition.

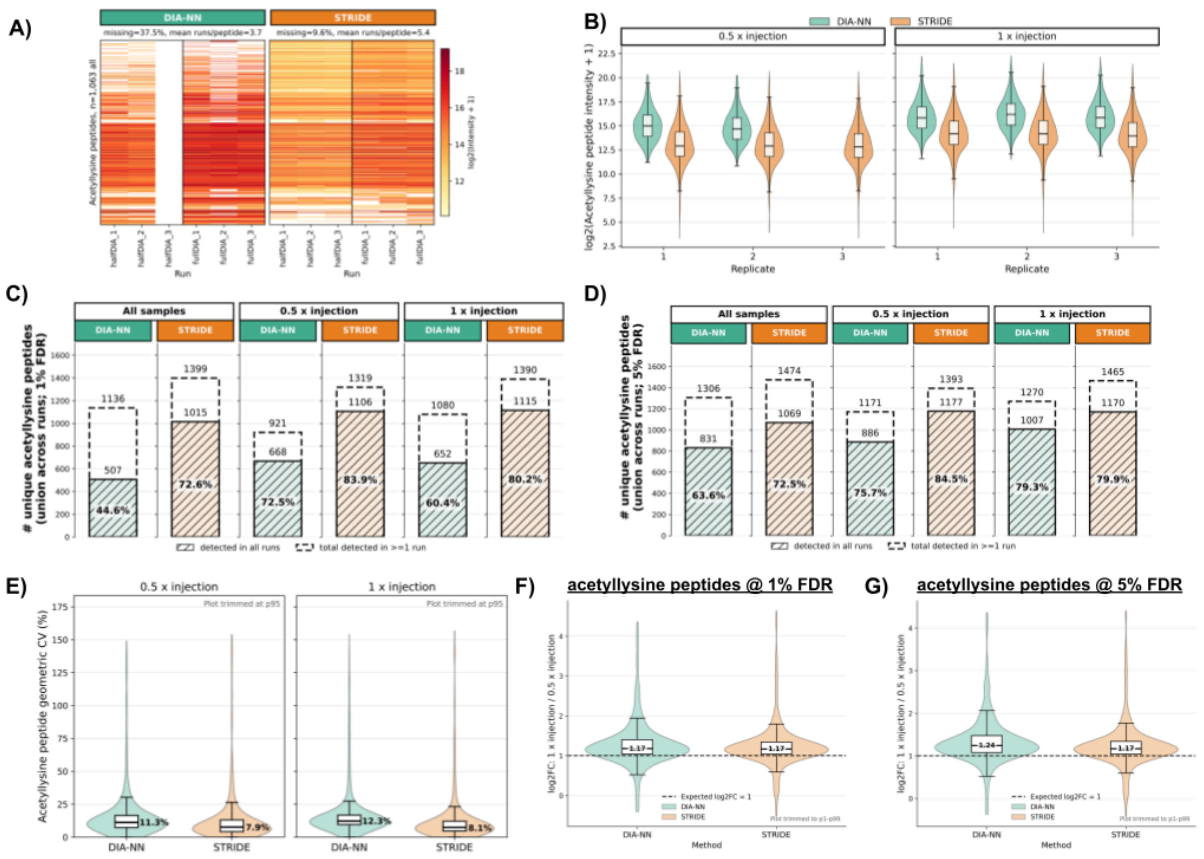


**Supplementary Figure 12. STRIDE improves acetyllysine quantification completeness and preserves expected intensity scaling. A)** Quantification heatmaps for the intersecting set of acetyllysine peptides detected by DIA-NN and STRIDE. White cells indicate missing quantification values. **B)** Distribution of log2 acetyllysine peptide intensities across replicate runs for each injection condition. **C-D)** Complete versus total acetyllysine peptide identifications at 1% FDR (**C**) and 5% FDR (**D**). Dashed bars show the total number of unique acetyllysine peptides detected in at least one run, while hatched bars show the subset detected in all replicate runs. **E)** Distribution of geometric coefficient of variation values for acetyllysine peptide quantification across replicate runs, shown separately for the two injection conditions. **F-G)** Distribution of log2 fold-changes between the 1x and 0.5x injection conditions for acetyllysine peptides identified at 1% FDR (**F**) and 5% FDR (**G**). The dashed horizontal line indicates the expected log2 fold-change of 1.

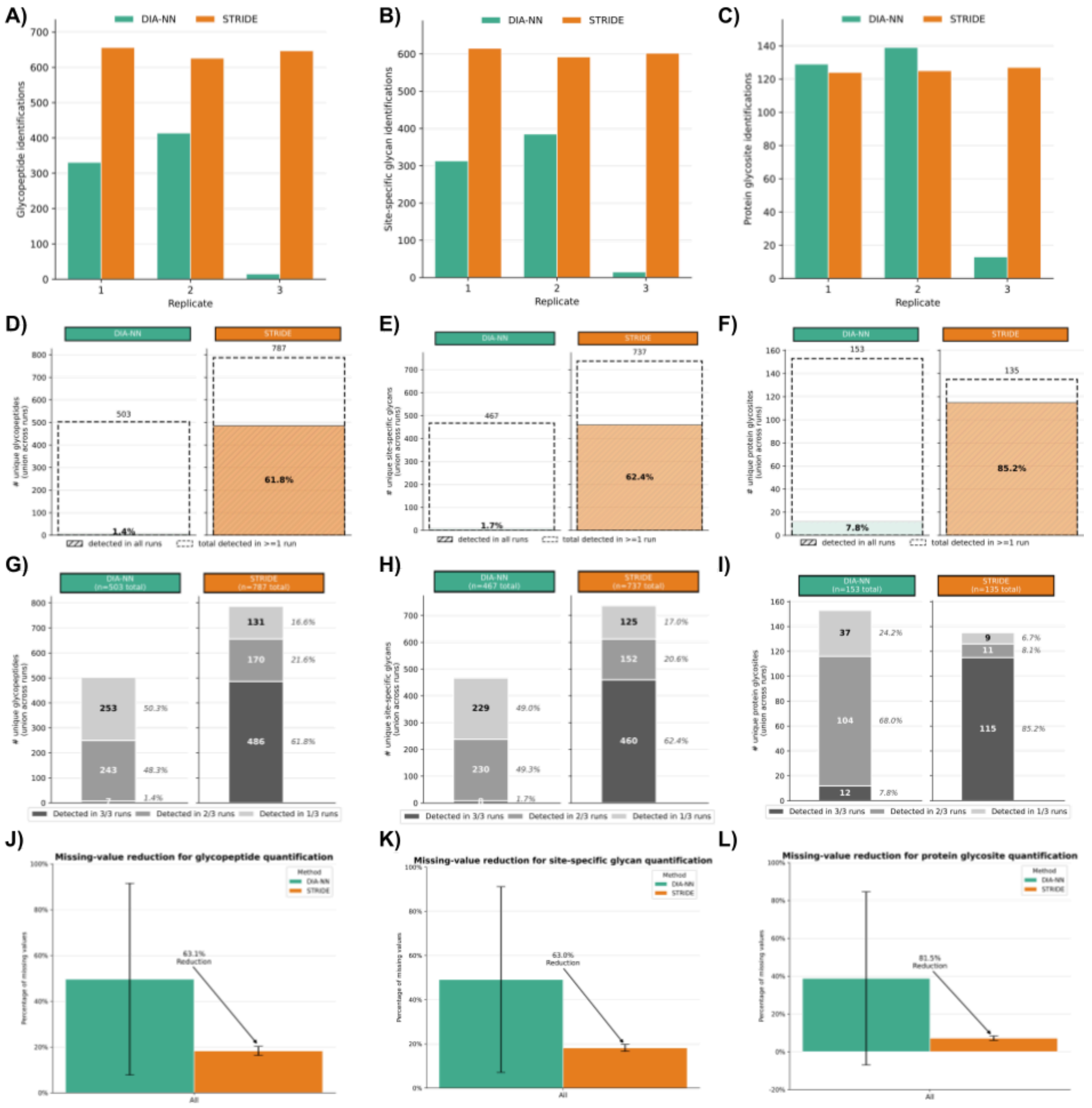


**Supplementary Figure 13. STRIDE improves identification consistency for human serum glycopeptides and glycoforms. A-C)** Number of identifications per replicate for glycopeptides (**A**), site-specific glycans (**B**), and protein glycosites (**C**). **D-F**) Complete versus total identifications for glycopeptides (**D**), site-specific glycans (**E**), and protein glycosites (**F**). Dashed bars show the total number of unique identifications detected in at least one run, while hatched bars show the subset detected in all three replicate runs. **G-I)** Reproducibility of identifications across replicate runs for glycopeptides (**G**), site-specific glycans (**H**), and protein glycosites (**I**). Stacked bars show the number and percentage of identifications detected in 3/3, 2/3, or 1/3 replicate runs. **J-L)** Mean percentage of missing quantification values across replicate runs for glycopeptides (**J**), site-specific glycans (**K**), and protein glycosites (**L**). Error bars show variation across runs, and arrows indicate the reduction in missing values obtained with STRIDE relative to DIA-NN.

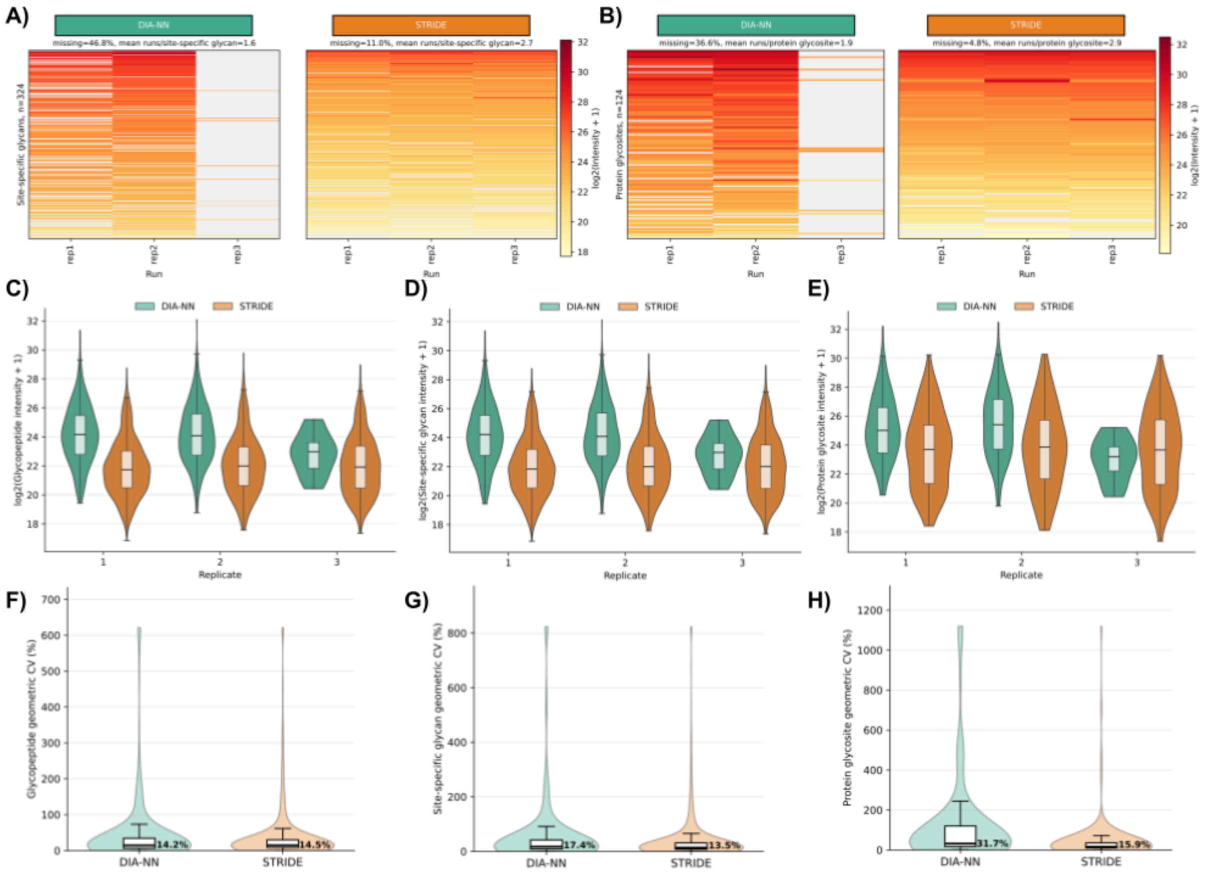


**Supplementary Figure 14. STRIDE improves quantification completeness for human serum glycopeptides and glycoforms. A-B)** Quantification heatmaps for site-specific glycans (**A**) and protein glycosites (**B**) across the three human serum replicate runs. White cells indicate missing quantification values. **C-E)** Distribution of log2 intensities across replicate runs for glycopeptides (**C**), site-specific glycans (**D**), and protein glycosites (**E**). **F-H)** Distribution of geometric CV values for glycopeptide (**F**), site-specific glycan (**Gf**), and protein glycosite (**H**) quantification. Median CV values are shown within each violin plot.

### STRIDE: Signal Transfer and Aligned Ion-Peak Discrimination for Peptidoform Evidence for Consistent Quantification of Site-Localized Post-Translational Modifications in Large-Scale DIA-MS: Supplementary Notes

*Justin Cyril Sing[1,2], Shubham Gupta[4], Hannes Luc Röst*[1,2,3]*

1. Donnelly Centre for Cellular and Biomolecular Research, University of Toronto, Toronto, Canada
2. Department of Molecular Genetics, University of Toronto, Toronto, Canada
3. Department of Computer Science, University of Toronto, Toronto, Canada
4. Department of Genetics, Stanford University, California, United States

### STRIDE Algorithm Overview

### Alignment and Cross Run Peak-Mapping

ARYCAL performs confidence-aware peak-group alignment in four successive stages:

1. from detecting-transition XICs to run-specific TIC traces,
2. from TIC traces to reference-aligned RT mappings using FFT-initialized DTW,
3. from reference peak-groups to projected query-run target locations, and
4. from query-run candidate peak-groups to one-to-one mappings with direct mapping confidence.

The resulting framework combines chromatogram-based RT alignment with confidence-aware across-run peak-group mapping for downstream signal propagation, such as peptidoform inference.

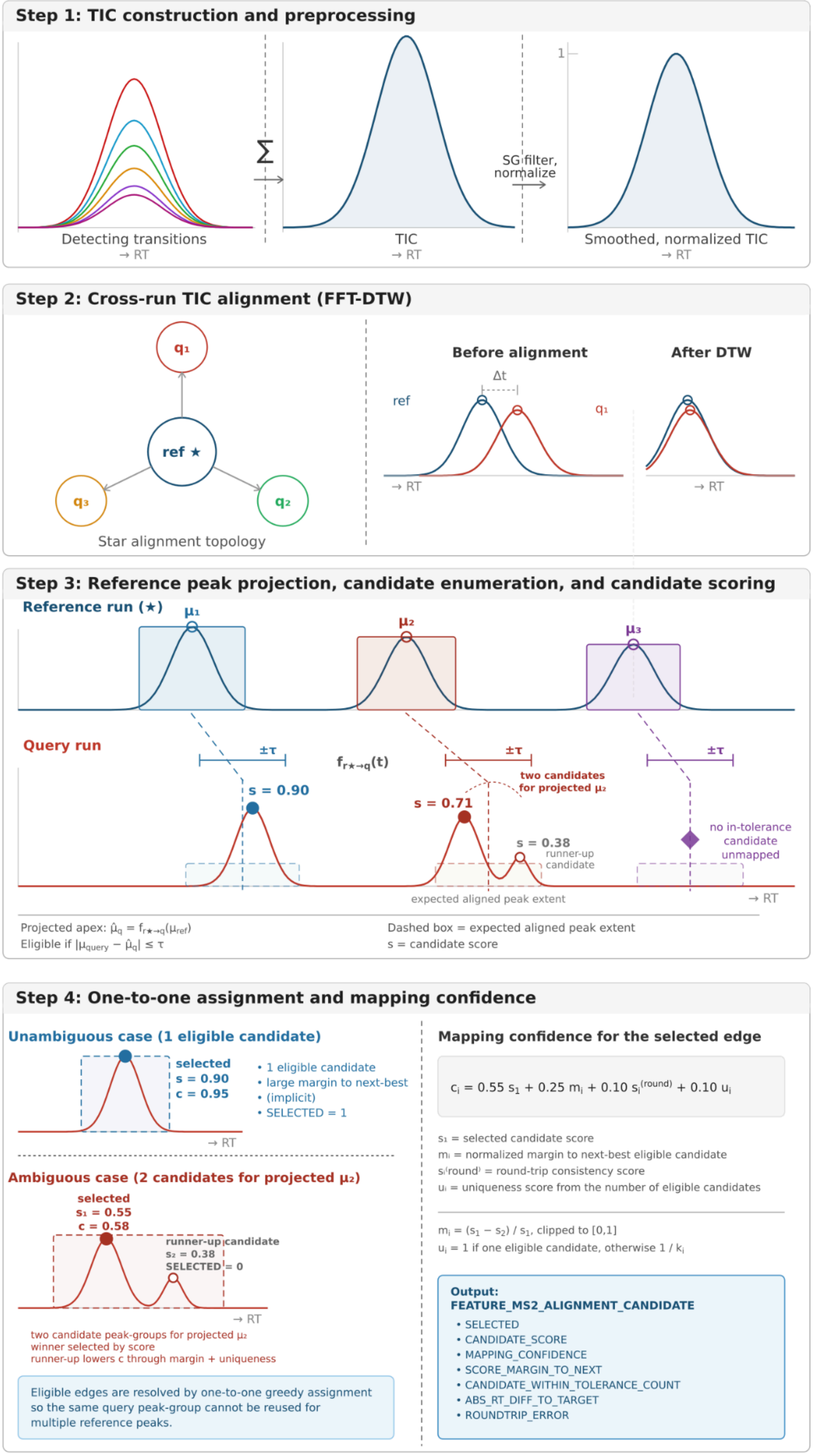

Step 1: TIC construction and preprocessing
Σ
SG filter, normalize
1
Detecting transitions
→ RT
TIC
→ RT
Smoothed, normalized TIC
→ RT
Step 2: Cross-run TIC alignment (FFT-DTW)
$q_1$
ref ★
$q_3$
$q_2$
Star alignment topology
Before alignment
After DTW
Δt
ref
$q_1$
→ RT
→ RT
Step 3: Reference peak projection, candidate enumeration, and candidate scoring
Reference run (★)
$\mu_1$
$\mu_2$
$\mu_3$
Query run
±τ
±τ
±τ
$f_{r\star\to q}(t)$
two candidates for projected $\mu_2$
s = 0.90
s = 0.71
s = 0.38
runner-up candidate
no in-tolerance candidate unmapped
expected aligned peak extent
→ RT
Projected apex: $\hat{\mu}_q = f_{r\star\to q}(\mu_{ref})$
Eligible if $|\mu_{query} - \hat{\mu}_q| \le \tau$
Dashed box = expected aligned peak extent
s = candidate score
Step 4: One-to-one assignment and mapping confidence
Unambiguous case (1 eligible candidate)
selected
s = 0.90
c = 0.95
• 1 eligible candidate
• large margin to next-best
• (implicit)
• SELECTED = 1
→ RT
Ambiguous case (2 candidates for projected $\mu_2$)
selected
$s_1$ = 0.55
c = 0.58
runner-up candidate
$s_2$ = 0.38
SELECTED = 0
→ RT
two candidate peak-groups for projected $\mu_2$
winner selected by score
runner-up lowers c through margin + uniqueness
Eligible edges are resolved by one-to-one greedy assignment so the same query peak-group cannot be reused for multiple reference peaks.
Mapping confidence for the selected edge
$c_i = 0.55\, s_1 + 0.25\, m_i + 0.10\, s_i^{(round)} + 0.10\, u_i$
$s_1$ = selected candidate score
$m_i$ = normalized margin to next-best eligible candidate
$s_i^{(round)}$ = round-trip consistency score
$u_i$ = uniqueness score from the number of eligible candidates
$m_i = (s_1 - s_2) / s_1$, clipped to [0,1]
$u_i$ = 1 if one eligible candidate, otherwise $1 / k_i$
Output:
FEATURE_MS2_ALIGNMENT_CANDIDATE
• SELECTED
• CANDIDATE_SCORE
• MAPPING_CONFIDENCE
• SCORE_MARGIN_TO_NEXT
• CANDIDATE_WITHIN_TOLERANCE_COUNT
• ABS_RT_DIFF_TO_TARGET
• ROUNDTRIP_ERROR

**Supplementary Figure1. Overview of the across run dynamic chromatogram alignment (ARYCAL) and confidence-aware peak-group mapping workflow.**
For each precursor, ARYCAL first extracts detecting-transition chromatograms in each run and collapses them into a total ion chromatogram (TIC), followed by Savitzky-Golay smoothing and intensity normalization **(Step 1)**. Smoothed run-level TICs are then aligned to a common reference run using a star-alignment strategy with FFT-initialized dynamic time warping (FFT-DTW), yielding a nonlinear retention-time mapping between the reference and each query run **(Step 2)**. Reference-run peak-groups are projected into each query run using the learned RT mapping, producing a projected apex $\hat{\mu}_q$ and an expected aligned peak extent; all same-precursor query peak-groups are enumerated and scored as candidate mappings, and candidates are considered eligible when their apex RT satisfies $|\mu_{\text{query}} - \hat{\mu}_q| \leq \tau$ **(Step 3)**. Candidate scores are computed from RT agreement, projected-versus-observed boundary overlap, peak-width similarity, and optional peak-group metadata (e.g. rank and q-value). Eligible candidate edges are then resolved by one-to-one greedy assignment so that the same query peak-group cannot be reused for multiple reference peaks. For each selected edge, a final mapping confidence is computed as a weighted combination of the selected candidate score, the normalized score margin to the next-best eligible candidate, round-trip RT consistency, and a uniqueness term reflecting the number of in-tolerance candidates **(Step 4)**. The resulting candidate-level output table, FEATURE_MS2_ALIGNMENT_CANDIDATE, stores all evaluated reference-query peak-group pairs together with the selected mapping and mapping-confidence metrics for downstream filtering during across-run signal propagation.

### 1. Input Representation and Peak-Group Universe

For each precursor, ARYCAL reads:

- the detecting transition identifiers from the *OSW* assay definition, and optional precursor and identifying transition identifiers,
- the corresponding fragment-ion chromatograms from the XIC files, and
- the candidate peak-groups already present in the *OSW* `FEATURE`/`FEATURE_MS2` tables for each run.

ARYCAL can utilize the precursor chromatograms and identifying transitions during alignment. However, the default only uses the detecting transitions for alignment, i.e. driven exclusively by the detecting fragment-ion traces. The main reason for this reasonable default behaviour is to save I/O for reading XIC data, and to reduce memory consumption required to hold ~100+ XIC traces per precursor in memory.

Let precursor $p$ have $M_p$ detecting transitions and be observed in $R$ runs. For run $r$, transition $m$, and retention time $t$, the raw chromatographic signal is denoted

$$x_{p,r,m}(t).$$

OpenSWATH candidate peak-groups are represented as per-run tuples

$$g_{p,r,j} = (\text{feature\_id}_{p,r,j},\ \mu_{p,r,j},\ \ell_{p,r,j},\ u_{p,r,j},\ I_{p,r,j},\ \text{rank}_{p,r,j},\ q_{p,r,j}),$$

where $\mu$ is the apex RT, $\ell$ and $u$ are the left and right boundaries, $I$ is the peak-group intensity, *rank* is the OpenSWATH peak-group rank when available, and $q$ is the peak-group q-value when available.

### 2. TIC Construction and Preprocessing

#### 2.1 Detecting-transition TICs

For each run, ARYCAL collapses the detecting-transition chromatograms into a TIC:

$$T_{p,r}(t) = \sum_{m=1}^{M_p} x_{p,r,m}(t).$$

This TIC is not intended as a quantitative replacement for the fragment traces. Rather, it is a high-signal summary trace used to estimate the global RT transformation between runs.

#### 2.2 Common RT grid

Before alignment, ARYCAL constructs a precursor-specific common RT grid by taking the union of all observed RT values across runs:

$$\mathcal{T}_p = \bigcup_{r=1}^{R} \{t_{p,r,1}, t_{p,r,2}, \ldots\}.$$

Each run-specific TIC is then embedded into this shared grid. In the current implementation, missing time points are **zero-filled** rather than interpolated. This is an important implementation detail: the aligned TICs therefore preserve the sparse/jagged structure of the underlying extracted traces instead of introducing interpolation smoothness.

#### 2.3 Savitzky-Golay smoothing

Each common-grid TIC is smoothed using a Savitzky-Golay filter. For a window of length $W$ and polynomial order $d$, the smoothed TIC value at index $i$ is obtained by fitting a local polynomial

$$P_i(z) = \sum_{k=0}^{d} a_{i,k} z^k$$

to the intensities inside the window centered at $i$, and evaluating $P_i$ at the center point. In the current implementation, the default is set to: $W = 11$ and $d = 3$. Negative smoothed intensities are truncated to zero.

##### 2.4 Min-max normalization

Each smoothed TIC is then min-max normalized:

$$m = \min_t T_{p,r}^{(\mathrm{sg})}(t), \qquad M = \max_t T_{p,r}^{(\mathrm{sg})}(t).$$

$$\tilde{T}_{p,r}(t) = \frac{T_{p,r}^{(\mathrm{sg})}(t) - m}{M - m}.$$

This normalization removes global scale differences between runs before cross-correlation and DTW.

#### 3. Across-Run TIC Alignment by FFT-DTW

##### 3.1 Star alignment topology

ARYCAL supports star, minimum-spanning-tree, and progressive alignment topologies. In the configuration described here, `reference_type = "star"` is used together with an explicit `reference_run`, so every run is aligned directly to a single fixed reference run. If no reference run is supplied, the current implementation falls back to selecting a run at random.

For a precursor $p$, denote the reference TIC as $\widetilde{T}_{p,r^\star}$ and a query TIC as $\widetilde{T}_{p,r}$.

##### 3.2 FFT cross-correlation

The first alignment stage estimates a coarse lag by full cross-correlation. ARYCAL computes the full correlation by FFT using the convolution identity

$$c_{r^\star,r}[\ell] = \sum_t \widetilde{T}_{p,r^\star}(t)\,\widetilde{T}_{p,r}(t-\ell),$$

implemented as

$$c = \widetilde{T}_{p,r^\star} * \mathrm{reverse}(\widetilde{T}_{p,r}).$$

The lag estimate is

$$\hat{\ell}_{r^\star,r} = \arg\max_{\ell} c_{r^\star,r}[\ell].$$

This lag is used only to initialize local refinement.

##### 3.3 DTW refinement

After shifting the query TIC by the FFT lag, ARYCAL performs local refinement with dynamic time warping. For reference indices $i$ and query indices $j$, the accumulated DTW cost satisfies the standard recurrence

$$D(i,j) = d(i,j) + \min\{D(i-1,j), D(i,j-1), D(i-1,j-1)\},$$

where $d(i,j)$ is the local distance between reference and lag-shifted query intensities. The optimal path

$$\pi = \{(i_1,j_1),(i_2,j_2),\dots,(i_L,j_L)\}$$

defines the refined RT mapping between the reference TIC and the **original** query RT axis.

##### 3.4 RT mapping

ARYCAL stores the path as ordered pairs of RT values:

$$(\mathrm{rt}_k^{(1)}, \mathrm{rt}_k^{(2)}), \quad k = 1,\dots,L,$$

where $\mathrm{rt}^{(1)}$ belongs to the reference run and $\mathrm{rt}^{(2)}$ belongs to the original query run. For any reference RT $t$, the mapped query RT is obtained by piecewise-linear interpolation over neighboring path points:

$$\Delta t = \mathrm{rt}_{k+1}^{(1)} - \mathrm{rt}_k^{(1)}, \qquad \Delta r = \mathrm{rt}_{k+1}^{(2)} - \mathrm{rt}_k^{(2)}.$$

$$f_{r^\star\to r}(t) = \mathrm{rt}_k^{(2)} + \frac{t - \mathrm{rt}_k^{(1)}}{\Delta t}\,\Delta r.$$

with edge clamping outside the observed path range.

#### 4. Reference Peak Projection Across Runs

For each reference-run candidate peak-group $g_{p,r^\star,i}$, ARYCAL projects its apex and boundaries into each query run:

$$\hat{\mu}_{p,r,i} = f_{r^\star\to r}(\mu_{p,r^\star,i}),$$

$$\hat{\ell}_{p,r,i} = f_{r^\star\to r}(\ell_{p,r^\star,i}), \qquad \hat{u}_{p,r,i} = f_{r^\star\to r}(u_{p,r^\star,i}).$$

The pair $(\hat{\ell}_{p,r,i}, \hat{u}_{p,r,i})$ defines the expected aligned window for that reference peak-group in the query run.

ARYCAL also computes a round-trip consistency error by mapping the projected RT back to the reference axis:

$$e^{(\mathrm{rt})}_{\mathrm{round},p,r,i} = \left| f^{-1}_{r\to r^\star}\left(\hat{\mu}_{p,r,i}\right) - \mu_{p,r^\star,i} \right|.$$

This term measures self-consistency of the learned RT transformation around the projected peak.

#### 5. Candidate Enumeration Within Each Query Run

For each projected reference peak $i$ in query run $r$, ARYCAL enumerates all candidate peak-groups $g_{p,r,j}$ already present for precursor $p$ in that run. Each candidate is compared to the projected target RT $\hat{\mu}_{p,r,i}$.

Define

$$\Delta^{(\mathrm{target})}_{i,j} = \left|\mu_{p,r,j} - \hat{\mu}_{p,r,i}\right|, \qquad \Delta^{(\mathrm{ref})}_{i,j} = \left|\mu_{p,r,j} - \mu_{p,r^\star,i}\right|.$$

A candidate is considered **within tolerance** if

$$\Delta_{i,j}^{(\text{target})} \leq \tau,$$

where $\tau$ is the user-defined RT tolerance (`rt_mapping_tolerance`, 25 s in the current configuration).

If at least one candidate lies within tolerance, only those candidates remain eligible for assignment. If no candidate lies within tolerance, ARYCAL retains only the nearest candidate for reporting and diagnostics, but it does not promote that fallback candidate into the selected one-to-one assignment.

### 6. Confidence-Aware Candidate Scoring

ARYCAL scores every reference-query candidate pair using a weighted combination of RT agreement, boundary agreement, and existing peak-group metadata.

#### 6.1 Component scores

For candidate $j$ relative to projected reference peak $i$:

1. **RT score**

$$s_{i,j}^{(\text{rt})} = \max\left(0,\ 1 - \frac{\Delta_{i,j}^{(\text{target})}}{\tau}\right).$$

2. **Width overlap score** (intersection-over-union of projected and observed windows)

Let

$$A_i = [\hat{\ell}_{p,r,i}, \hat{u}_{p,r,i}], \qquad B_j = [\ell_{p,r,j}, u_{p,r,j}].$$

Then

$$s_{i,j}^{(\text{ov})} = \frac{|A_i \cap B_j|}{|A_i \cup B_j|}.$$

3. **Width similarity score**

Let projected width $w_i = \max(1, |\hat{u}_{p,r,i} - \hat{\ell}_{p,r,i}|)$ and candidate width $w_j = \max(1, |u_{p,r,j} - \ell_{p,r,j}|)$. Then

$$s_{i,j}^{(\mathrm{width})} = \max\left(0,\ 1 - \frac{|w_j - w_i|}{\max(w_i, w_j)}\right).$$

4. **Rank score**

If the candidate carries an OpenSWATH peak-group rank $r_j$,

$$s_{i,j}^{(\mathrm{rank})} = \frac{1}{\max(r_j, 1)}.$$

If rank is unavailable, ARYCAL uses a neutral fallback of 0.5.

5. **q-value score**

If the candidate carries an OpenSWATH q-value $q_j$,

$$s_{i,j}^{(\mathrm{q})} = 1 - \min(\max(q_j, 0), 1).$$

If q-value is unavailable, ARYCAL again uses a neutral fallback of 0.5.

6. **Intensity similarity score**

If the reference and query intensities are both available,

$$s_{i,j}^{(\mathrm{int})} = \frac{1}{1 + |\log(I_{p,r,j} + 1) - \log(I_{p,r^\star,i} + 1)|}.$$

If either intensity is unavailable, the implementation uses a neutral fallback of 0.5.

#### 6.2 Candidate score

The final candidate score is the convex combination

$$s_{i,j} = 0.45\, s_{i,j}^{(\mathrm{rt})} + 0.20\, s_{i,j}^{(\mathrm{ov})} + 0.10\, s_{i,j}^{(\mathrm{width})} + 0.10\, s_{i,j}^{(\mathrm{rank})} + 0.10\, s_{i,j}^{(\mathrm{q})} + 0.05\, s_{i,j}^{(\mathrm{int})},$$

clamped to $[0, 1]$.

The feature design intentionally emphasizes RT agreement and projected-boundary overlap, while allowing pre-existing peak-group rank/q-value to regularize the decision when multiple nearby candidates are present.

#### 7. One-to-One Cross-Run Assignment

For each query run, ARYCAL does not independently choose the nearest candidate for every reference peak. Instead, it performs a one-to-one greedy assignment over all eligible reference-query edges within the precursor and run pair.

Eligible edges are sorted by:

1. descending candidate score $s_{i,j}$,
2. ascending $\Delta_{i,j}^{(\mathrm{target})}$,
3. ascending peak-group rank,
4. ascending feature identifier.

ARYCAL then traverses this ordered list and accepts an edge only if:

- the corresponding reference peak has not already been assigned, and
- the query feature has not already been used by another reference peak.

This produces a sparse, non-overlapping set of reference-query mappings. Operationally, this is a greedy maximum-score bipartite assignment rather than a strict nearest-neighbor RT matcher.

#### 8. Mapping Confidence

For each selected mapping, ARYCAL computes an additional confidence score designed specifically for downstream filtering.

Let $s_1$ be the selected candidate score and $s_2$ the best competing score among the remaining **within-tolerance** candidates for the same reference peak. The raw score margin is

$$\delta_i = \max(s_1 - s_2, 0).$$

If no second candidate exists, the implementation defaults to $\delta_i = s_1$. The normalized margin is

$$m_i = \begin{cases} \delta_i / s_1, & s_1 > 0, \\ 0, & s_1 = 0. \end{cases}$$

The RT round-trip score is

$$s_i^{(\text{round})} = \max\left(0,\ 1 - \frac{e^{(\text{rt})}_{\text{round},i}}{\tau}\right).$$

The uniqueness score is

$$u_i = \begin{cases} 1, & k_i \le 1, \\ 1/k_i, & k_i > 1, \end{cases}$$

where $k_i$ is the number of within-tolerance candidates for reference peak $i$.

The final mapping confidence is

$$c_i = 0.55\, s_1 + 0.25\, m_i + 0.10\, s_i^{(\text{round})} + 0.10\, u_i,$$

again clamped to $[0,1]$.

This confidence has a clear interpretation:

- high when the selected candidate is intrinsically strong,
- high when it clearly outranks nearby alternatives,
- high when the local RT mapping is self-consistent, and
- high when the local candidate space is unambiguous.

### 9. Output Tables

#### 9.1 Candidate confidence table

`FEATURE_MS2_ALIGNMENT_CANDIDATE` contains **all** candidate mappings evaluated for each reference peak in each query run, including:

- projected target RT (`MAPPED_TARGET_RT`)
- expected mapped boundaries
- whether the candidate was selected
- candidate score
- mapping confidence (for selected candidates)
- score margin to next-best candidate
- normalized RT error
- absolute RT differences to projected and reference RT
- RT round-trip error
- width, rank, q-value, and intensity-based sub-scores

This table is the primary output for confidence-aware propagation. In the current OpenMS and local PyProphet downstream adaptations, across-run signal propagation can be restricted to rows with

$$\mathrm{SELECTED} = 1 \quad \text{and} \quad \mathrm{MAPPING\_CONFIDENCE} \geq c_{\min},$$

where $c_{\min}$ is user-defined (e.g. 0.5-0.8 depending on the desired precision-recall trade-off).

### Bayesian Signal Propagation for Peptidoform Inference Across Runs

The algorithm propagates confidence in three successive stages:

1. from peak-group confidence to precursor-level confidence,
2. from precursor-level confidence to peptidoform-level posterior probabilities, and
3. optionally across runs by importing already-confident transition evidence between aligned peak groups.

The resulting framework combines within-run evidence integration with confidence-aware across-run evidence transfer.

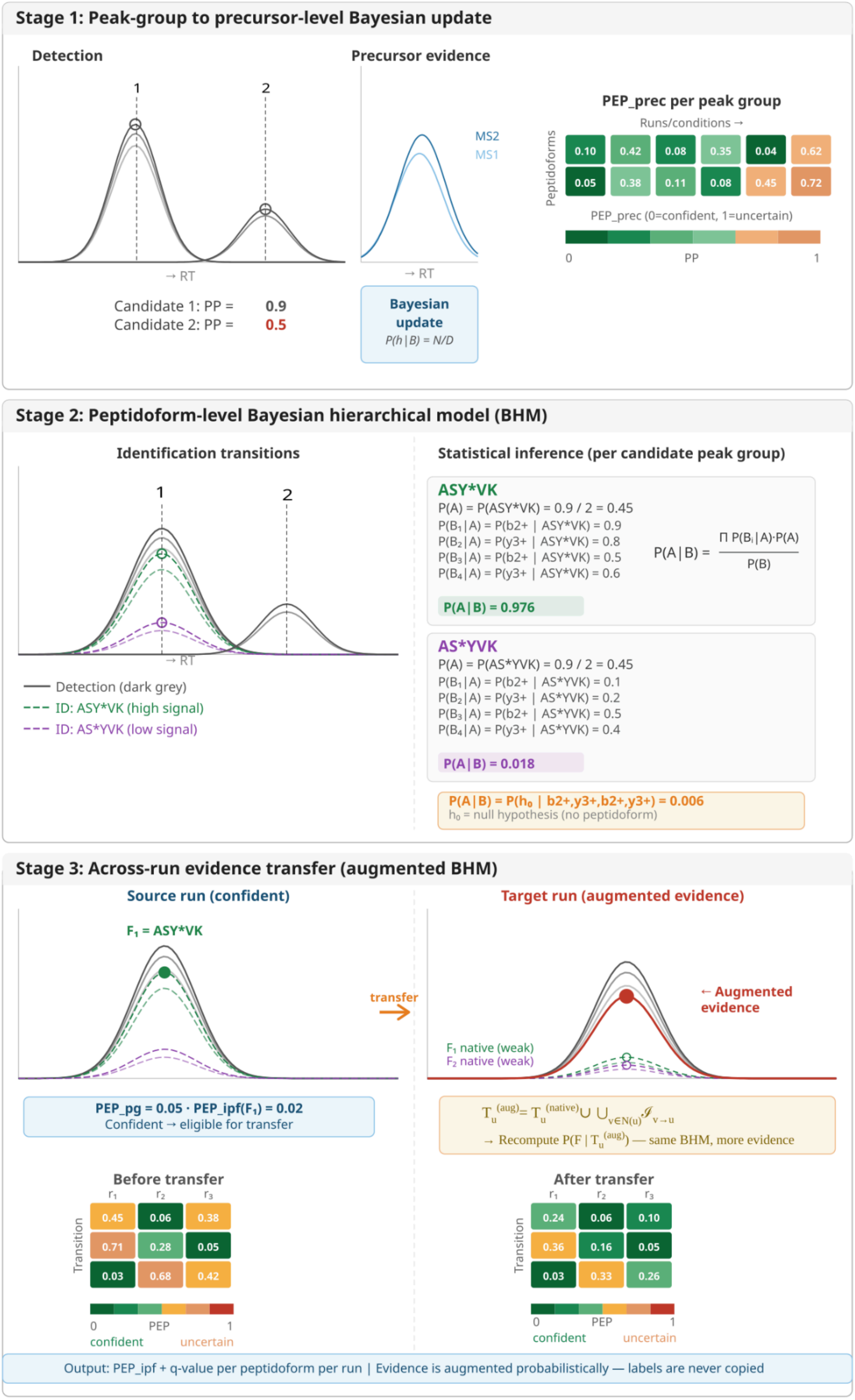

Stage 1: Peak-group to precursor-level Bayesian update
Detection
1
2
→ RT
Candidate 1: PP = 0.9
Candidate 2: PP = 0.5
Precursor evidence
MS2
MS1
→ RT
Bayesian update
P(h | B) = N/D
PEP_prec per peak group
Runs/conditions →
Peptidoforms
0.10 0.42 0.08 0.35 0.04 0.62
0.05 0.38 0.11 0.08 0.45 0.72
PEP_prec (0=confident, 1=uncertain)
0
PP
1
Stage 2: Peptidoform-level Bayesian hierarchical model (BHM)
Identification transitions
1
2
→ RT
Detection (dark grey)
ID: ASY*VK (high signal)
ID: AS*YVK (low signal)
Statistical inference (per candidate peak group)
ASY*VK
P(A) = P(ASY*VK) = 0.9 / 2 = 0.45
$P(B_1 | A)$ = P(b2+ | ASY*VK) = 0.9
$P(B_2 | A)$ = P(y3+ | ASY*VK) = 0.8
$P(B_3 | A)$ = P(b2+ | ASY*VK) = 0.5
$P(B_4 | A)$ = P(y3+ | ASY*VK) = 0.6
$P(A | B) = \frac{\prod P(B_i | A) \cdot P(A)}{P(B)}$
P(A | B) = 0.976
AS*YVK
P(A) = P(AS*YVK) = 0.9 / 2 = 0.45
$P(B_1 | A)$ = P(b2+ | AS*YVK) = 0.1
$P(B_2 | A)$ = P(y3+ | AS*YVK) = 0.2
$P(B_3 | A)$ = P(b2+ | AS*YVK) = 0.5
$P(B_4 | A)$ = P(y3+ | AS*YVK) = 0.4
P(A | B) = 0.018
$P(A | B) = P(h_0$ | b2+,y3+,b2+,y3+) = 0.006
$h_0$ = null hypothesis (no peptidoform)
Stage 3: Across-run evidence transfer (augmented BHM)
Source run (confident)
$F_1$ = ASY*VK
transfer
$PEP\_pg = 0.05 \cdot PEP\_ipf(F_1) = 0.02$
Confident → eligible for transfer
Target run (augmented evidence)
← Augmented evidence
$F_1$ native (weak)
$F_2$ native (weak)
$T_u^{(aug)} = T_u^{(native)} \cup \bigcup_{v \in N(u)} \mathcal{I}_{v \to u}$
→ Recompute $P(F | T_u^{(aug)})$ — same BHM, more evidence
Before transfer
$r_1$ $r_2$ $r_3$
Transition
0.45 0.06 0.38
0.71 0.28 0.05
0.03 0.68 0.42
0 PEP 1
confident uncertain
After transfer
$r_1$ $r_2$ $r_3$
Transition
0.24 0.06 0.10
0.36 0.16 0.05
0.03 0.33 0.26
0 PEP 1
confident uncertain
Output: PEP_ipf + q-value per peptidoform per run | Evidence is augmented probabilistically — labels are never copied

**Supplementary Figure2. Overview of the IPF signal propagation framework for across-run peptidoform inference.** The IPF workflow proceeds in three stages. In **Stage 1**, detection-level precursor evidence from MS1 and MS2 peak-group signals is combined within each run and candidate peak-group using a Bayesian update to produce a precursor-level posterior error probability ($\mathrm{PEP}_{\mathrm{prec}}$), which summarizes how confidently a given peak-group supports the presence of the precursor. In **Stage 2**, identification transitions are evaluated within each candidate peak-group using a Bayesian hierarchical model (BHM) to infer peptidoform identity from the observed fragment-ion evidence. This yields posterior probabilities and peptidoform-specific error estimates ($\mathrm{PEP}_{\mathrm{ipf}}$) for each candidate peak-group, while retaining an explicit null hypothesis when no peptidoform is sufficiently supported. In **Stage 3**, across-run signal propagation augments the evidence available in weak target runs using aligned, high-confidence peptidoform evidence from source runs. Rather than copying discrete labels across runs, the method transfers probabilistic evidence into the target run and recomputes the same hierarchical Bayesian model using the augmented evidence set. As a result, native weak signals can be reinforced by concordant across-run evidence, leading to improved peptidoform confidence and downstream q-values, while preserving uncertainty and allowing unsupported hypotheses to remain unassigned.

### 1. Problem Setup

Consider a set of runs $r \in \mathcal{R}$. For each precursor in each run, candidate peak groups are detected. A candidate peak group $g$ is associated with:

- a peak-group posterior error probability $p_g^{(\mathrm{pg})}$,
- optional precursor-specific evidence on one or more precursor channels,
- a set of candidate peptidoforms $\mathcal{F}_g = \{F_1, \dots, F_{N_g}\}$,
- a set of scored identification transitions,
- and optionally a set of cross-run alignments linking $g$ to corresponding peak groups in other runs.

The task is to estimate, for every candidate peptidoform $F \in \mathcal{F}_g$, the posterior probability that $F$ explains the signal observed in $g$.

### 2. Notation

For a candidate peak group $g$:

- $p_g^{(\mathrm{pg})}$ is the peak-group posterior error probability.
- $B_g$ is the set of retained precursor evidence signals for $g$.
- $p_b$ is the posterior error probability of precursor evidence item $b \in B_g$.
- $\mathcal{F}_g = \{F_1, \dots, F_{N_g}\}$ is the set of candidate peptidoforms for $g$.

- $N_g$ is the number of candidate peptidoforms.
- $\mathcal{T}_g$ is the set of retained transition evidence rows for $g$.
- $p_t$ is the posterior error probability of transition evidence row $t \in \mathcal{T}_g$.
- $s_{t,F} \in \{0, 1\}$ is the support indicator for transition $t$ with respect to peptidoform $F$.
- $h_0$ is the null hypothesis that the observed signal does not support any candidate peptidoform.

Two confidence thresholds are used for across-run propagation:

- $\tau_{\text{map}}$ for accepting cross-run peak mappings,
- $\tau_{\text{prop}}$ for deciding whether a transition row is confident enough to be transferred across runs.

### 3. Algorithm Overview

The algorithm can be summarized as follows:

1. detect candidate peak groups and assign peak-group posterior error probabilities;
2. refine each peak group to precursor level by combining the peak-group prior with precursor-specific evidence;
3. expand transition evidence against all candidate peptidoform hypotheses and evaluate a peptidoform-level Bayesian model;
4. if cross-run mappings are available, transfer only already-confident transition evidence across aligned peak groups and recompute the same peptidoform-level Bayesian model;
5. convert posterior error probabilities into local and global confidence metrics.

### 4. Precursor-Level Bayesian Update

#### 4.1 Hypothesis space

At precursor level, the signal is modeled with the binary hypothesis set

$$A_g^{(\text{prec})} = \{h, h_0\},$$

where $h$ is the true precursor state and $h_0$ is the false precursor state.

#### 4.2 Priors from peak-group confidence

The peak-group confidence defines the prior mass:

$$P(h) = 1 - p_g^{(\mathrm{pg})}, \qquad P(h_0) = p_g^{(\mathrm{pg})}.$$

Thus, the precursor model does not start from an uninformative prior. Instead, it uses the already-estimated confidence of the peak group as the starting point.

#### 4.3 Conditional evidence model

For every retained precursor evidence item $b \in B_g$, with error probability $p_b$,

$$P(b \mid h) = 1 - p_b, \qquad P(b \mid h_0) = p_b.$$

The precursor evidence sources are treated as conditionally independent:

$$P(B_g \mid a) = \prod_{b \in B_g} P(b \mid a), \qquad a \in \{h, h_0\}.$$

#### 4.4 Posterior update

The marginal likelihood is

$$P(B_g) = \sum_{a \in \{h, h_0\}} P(a) \prod_{b \in B_g} P(b \mid a).$$

The posterior probability of the true precursor state becomes

$$N = P(h) \prod_{b \in B_g} (1 - p_b), \qquad D = \sum_{a \in \{h, h_0\}} P(a) \prod_{b \in B_g} P(b \mid a).$$

$$P(h \mid B_g) = \frac{N}{D}.$$

The precursor-level posterior error probability is then

$$\mathrm{PEP}_g^{(\mathrm{prec})} = 1 - P(h \mid B_g).$$

##### 4.5 Interpretation

This stage shrinks or expands the error mass associated with the candidate peak group:

- if precursor evidence agrees with the peak group, $\mathrm{PEP}_g^{(\mathrm{prec})}$ decreases;
- if precursor evidence contradicts it, $\mathrm{PEP}_g^{(\mathrm{prec})}$ increases;
- if no precursor-specific evidence is available, the posterior reduces to the peak-group prior.

#### 5. Peptidoform-Level Bayesian Update

##### 5.1 Hypothesis set

For each candidate peak group $g$, the peptidoform model considers

$$A_g^{(\mathrm{ipf})} = \mathcal{F}_g \cup \{h_0\}.$$

The null hypothesis $h_0$ represents the possibility that the observed signal is not convincingly explained by any of the candidate peptidoforms.

##### 5.2 Prior redistribution

The precursor-level error mass is propagated directly to the null hypothesis:

$$P(h_0) = \mathrm{PEP}_g^{(\mathrm{prec})}.$$

The remaining confidence is distributed across the candidate peptidoforms:

$$P(F) = \frac{1 - \mathrm{PEP}_g^{(\mathrm{prec})}}{N_g}, \qquad F \in \mathcal{F}_g.$$

This construction ensures that

$$P(h_0) + \sum_{F \in \mathcal{F}_g} P(F) = 1.$$

#### 5.3 Transition support model

Each transition row is evaluated against every candidate peptidoform hypothesis, not only against the peptidoform it supports directly. The support indicator

$$s_{t,F} = \begin{cases} 1 & \text{if transition } t \text{ supports peptidoform } F \\ 0 & \text{otherwise} \end{cases}$$

controls whether the transition acts as positive or negative evidence.

For a transition row $t$, with posterior error probability $p_t$,

$$P(t \mid F) = \begin{cases} 1 - p_t & \text{if } s_{t,F} = 1 \\ p_t & \text{if } s_{t,F} = 0. \end{cases}$$

For the null hypothesis,

$$P(t \mid h_0) = p_t.$$

Thus, a transition that strongly supports one peptidoform automatically becomes strong opposing evidence for incompatible alternatives.

#### 5.4 Posterior update

Assuming conditional independence among retained transition rows,

$$P(\mathcal{T}_g \mid A) = \prod_{t \in \mathcal{T}_g} P(t \mid A).$$

The marginal is

$$P(\mathcal{T}_g) = \sum_{A \in A_g^{(\mathrm{ipf})}} P(A) \prod_{t \in \mathcal{T}_g} P(t \mid A).$$

The posterior probability of peptidoform $F$ is

$$N = P(F) \prod_{t \in \mathcal{T}_g} P(t \mid F), \qquad D = \sum_{A \in A_g^{(\mathrm{ipf})}} P(A) \prod_{t \in \mathcal{T}_g} P(t \mid A).$$

$$P(F \mid \mathcal{T}_g) = \frac{N}{D}.$$

The corresponding posterior error probability is

$$\mathrm{PEP}^{(\mathrm{ipf})}_{g,F} = 1 - P(F \mid \mathcal{T}_g).$$

##### 5.5 Why shared transitions still help

The transition model is not limited to uniquely diagnostic transitions. A shared transition can still contribute evidence because it supports multiple compatible hypotheses while simultaneously opposing incompatible ones. The method therefore captures both:

- **discriminative evidence** from peptidoform-specific transitions,
- **contextual evidence** from shared transitions that remain informative in the presence of the full candidate set.

This is especially important when only a small number of uniquely diagnostic transitions are observable.

#### 6. Across-Run Evidence Transfer

##### 6.1 Cross-run peak mapping

Across-run propagation begins with a peak-mapping stage that links candidate peak groups across runs. For a given precursor context, the result is a graph of aligned peak groups. Let

$$G = (\mathcal{V}, \mathcal{E})$$

be the alignment graph, where vertices are candidate peak groups and an edge $(u, v)$ is retained only if its mapping confidence

$$m_{u,v} \geq \tau_{\mathrm{map}}.$$

Each connected component defines a set of peak groups that are eligible to exchange evidence.

#### 6.2 Native evidence and transferred evidence

For a target peak group $u$, let

$$\mathcal{T}_u^{(\text{native})}$$

denote its native transition evidence.

From every aligned source peak group $v$ in the same connected component, only already-confident transition rows are imported:

$$\mathcal{I}_{v \to u} = \left\{ t \in \mathcal{T}_v^{(\text{native})} \;\middle|\; p_t \le \tau_{\text{prop}} \right\}.$$

The augmented evidence set for the target peak group is

$$\mathcal{T}_u^{(\text{aug})} = \mathcal{T}_u^{(\text{native})} \cup \bigcup_{v \in \mathcal{N}(u)} \mathcal{I}_{v \to u},$$

where $\mathcal{N}(u)$ denotes aligned neighbors of $u$.

Two properties are essential:

1. native evidence is always retained;
2. transferred evidence is imported only if it is already locally confident in the source run.

#### 6.3 Duplicate collapse

After evidence transfer, the same transition-hypothesis pair may be represented more than once in the target evidence set. These duplicates are collapsed before the Bayesian update.

Conceptually, for duplicated instances of the same transition-hypothesis relation, the transferred evidence is summarized by the most confident available instance:

$$p_t^{(\text{collapsed})} = \min_i p_t^{(i)}.$$

This avoids overweighting repeated representations of the same transition assignment while preserving the strongest available support.

##### 6.4 Posterior recomputation rather than label copying

Across-run propagation does not impose a peptidoform label from one run onto another. Instead, it augments the evidence set of the target peak group and then recomputes the same Bayesian peptidoform posterior:

$$P(F \mid \mathcal{T}_u^{(\mathrm{aug})}).$$

The transferred evidence therefore acts as additional probabilistic support, not as a deterministic override.

#### 7. Error-Rate Interpretation

The posterior error probability of a peptidoform hypothesis,

$$\mathrm{PEP}_{g,F}^{(\mathrm{ipf})},$$

can be interpreted as a local error estimate.

To obtain a global error estimate across a family of peptidoform calls, candidates are sorted by increasing posterior error probability:

$$\mathrm{PEP}_{(1)} \le \mathrm{PEP}_{(2)} \le \cdots \le \mathrm{PEP}_{(M)}.$$

The corresponding monotone global error estimate can be written as

$$q_{(k)} = \min_{\ell \ge k} \frac{1}{\ell} \sum_{i=1}^{\ell} \mathrm{PEP}_{(i)}.$$

If desired, this operation can be applied:

- globally across all hypotheses, or
- separately within strata defined by the number of candidate peptidoforms $N_g$.

The second strategy is useful when highly ambiguous precursor contexts should be benchmarked primarily against similarly ambiguous cases.
